# Insights into the Microbial Weathering of Chinese Wooden Ancestral Halls in Guangdong Province

**DOI:** 10.1101/2023.01.19.524717

**Authors:** Manchun Liu, Xining Su, Qinqing Wen, Tongshu Yang, Muyuyang Lin, Paierzhati Abudureyimu, Jerome Rumdon Lon, Jianfei Luo

**Affiliations:** School of Biology and Biological Engineering, South China University of Technology, Guangzhou, China; Institute of Synthetic Biology, Shenzhen Institute of Advanced Technology, Shenzhen, China

**Keywords:** wooden ancestral hall, biodeterioration, amplicon sequencing, environmental factors, biocides

## Abstract

Wooden buildings are facing biodeterioration due to natural weathering. Ancestral halls are traditional Chinese wooden architecture with high artistic value. The research analyzed the microbial diversity of nine ancestral halls in Guangdong Province under subtropical monsoon climates. With amplicon sequencing, we found a common core microorganism group, including *Bacillus*, *Pseudomonas*, *Paenibacillus*, *Acinetobacter*, *Toxicocladosporium*, *Cladosporium*, *Aspergillus*, and *Epicoccum*. A total of 8 different species of bacteria and 7 different species of fungi were isolated from Chen Clan Ancestral halls, most of which were proven to be capable of degrading cellulose or lignin. The comparison between damaged and undamaged sites (whether the paint coating is flaked off) revealed similar microbial compositions but with differing abundances. *Firmicutes* and *Ascomycota* dominated damaged surfaces, while *Proteobacteria* and *Actinobacteria* prevailed in undamaged ones. Notably, *Cladosporium*, widely present in both sites, emerged as the primary cellulose-degrading genus. Given the similar microbial compositions between damaged and undamaged sites, it is suggested that using paint alone as the exclusive means to protect bare wood is insufficient. Apparent correlation between microorganisms on wooden surfaces and humans showed that the increase of visitor flow rates might impact the microbiota composition on cultural relics. We evaluated the effect of four representative biocides and determined the feasibility of low-concentration Isothiazolinone. This research facilitates a basic understanding of the microbial weathering of ancestral halls and add significant reference for preserving wooden structures.

## 1. Introduction

Bare wooden architecture in open-air conditions provides a common habitat for numerous microorganisms, including bacteria and fungi, owing to its porous structure and the organic components (Cennamo et al., 2018; De Windt et al., 2014). Constituting approximately 40-60% cellulose, 15-30% hemicellulose, and 17-35% lignin(Velasco-Rodriguez et al., 2022), wood offers a rich nutritional source for microbial colonization. Microbes then form biofilms on the wooden surfaces, developing an extracellular polysaccharide matrix (EPS) comprised of cell debris, proteins, polysaccharides, and lipids. These biofilms create a protective microenvironment fostering microbial growth, thereby hastening the degradation of wooden structures. (Coelho et al., 2021; Li et al., 2022; Negi & Sarethy, 2019; Skipper et al., 2022). Moreover, the pigments produced by microorganisms, along with the diverse colors exhibited by the microbial communities themselves, blanket the surface of wooden cultural relics, resulting in significant aesthetic damage (Kim et al., 2020; Sterflinger & Pinar, 2013). Additionally, Cellulolytic microorganisms such as *Bacillus, Aspergillus, Cladosporium*, and *Penicillium* have been identified for their capability to degrade Polychrome wood artifacts.

(Cennamo et al., 2018). Furthermore, enzymes such as cellulase, hemicellulase, and laccase, released during microbial metabolic processes, actively deteriorate cellulose and lignin, impacting the structural integrity of wooden cultural relics (Huang et al., 2021; Zhang et al., 2021). Therefore, research into microbial weathering causing biodeterioration in wooden architecture is essential. Ortiz et al. collected wood samples from eight historic churches in Chiloé, identifying fungi species such as *Coniophora, Laetiporus,* and *Postia* (causing brown rot), as well as *Trichoderma, Alternaria,* and *Verticillium* (associated with wood decay) at various stages of decay(Ortiz et al., 2014). Wang et al. utilized high-throughput sequencing to identify *Bacteroidetes, Proteobacteria, Firmicutes,* and *Actinobacteria* as the primary sources of bacterial cellulose-decomposing enzymes in an ancient wooden seawall in the Qiantang River of Hangzhou and they also isolated 11 cellulose-decomposing strains (Wang et al., 2023).

Ancestral halls, traditional Chinese wooden architecture prevalent in southern China, date back to the Ming (AD1368 to AD1644) and Qing (AD1636 to AD1912) Dynasties. These structures were initially erected to honor ancestors and preserve family legacies, holding significant educational and commemorative value for clans. Their intricate architecture, deep historical significance, and cultural resonance have had a lasting global influence. Despite their importance, many ancestral halls have suffered structural and aesthetic damage due to prolonged exposure to microbial, physical, and chemical weathering. Notably, limited research exists on the microbial weathering affecting these invaluable structures.

Currently, wooden ancestral halls are often coated with paint for microbial inhibition(Gobakken et al., 2010), with urushiol commonly included for its antimicrobial properties(Kim et al., 2021).

However, studies reveal the temporary nature of this safeguard. Research on eight hardwood surfaces treated with commonly used acrylic paint and exposed to open air demonstrates degradation within a year, with even the most durable coating showing signs of cracking and peeling within five years.(De Windt et al., 2014). Factors like increased humidity, mechanical stress, and physical weathering contribute to coating swelling, mottling, and eventual flaking, exposing the wooden surface.(Phulpoto et al., 2020). Hence, discussions persist regarding the effectiveness and limitations of using paint coatings for antimicrobial preservation on ancestral halls. Therefore, it is crucial to compare the microbial compositions between damaged and undamaged sites (whether the paint coating is flaked off).

Environmental factors can affect the diversity of microbiota on wooden cultural relics. For instance, human activities, even in the absence of direct interaction, contribute to about 25 to 30% of human skin-associated microbial taxa found on built environment surfaces (Gibbons, 2016). Increased tourism introduces skin or hair-borne microorganisms that can disrupt the existing microbial equilibrium on cultural relics (Pasquarella et al., 2015). Additionally, Zhang et al. highlighted the correlation between environmental factors such as temperature, humidity, and pH with variations in microbial communities on the Sandstone of Beishiku Temple (Zhang et al., 2023).

Merely applying a coating of paint does not entirely prevent microbial weathering on wooden relics. Microorganisms known for causing biodeterioration and biodegradation are commonly found on painted surfaces, including bacteria like *Bacillus*, *Pseudomonas*, *Enterobacter*, *Actinomycetes*, and fungi such as *Aspergillus*, *Trichoderma*, *Altemaria*, *Cladosporium*, and *Acremonium*(Hyvarinen et al., 2002; Phulpoto et al., 2020). Beyond paint coatings, the use of biocides is a prevalent method to combat microorganisms on wooden cultural relics. These biocides, comprising organic compounds, inorganic substances, and plant essential oils, present varying degrees of effectiveness(Kakakhel et al., 2019). While commercial biocides like Isothiazolinone, benzalkonium chloride, and sodium hypochlorite exhibit potent bacteriostatic effects, their high concentrations can pose environmental and human health risks(Kampf, 2018; Kozirog et al., 2016; Silva et al., 2020). Plant essential oils, lauded for their eco-friendliness, warrant further investigation regarding their antibacterial efficacy and practical application (Antonelli et al., 2020; Palla et al., 2020).

Given the significance of understanding microbial weathering mechanisms, it is important to classify microbial communities and elucidate the biological degradation process of cultural relics. To ensure uniform climatic conditions, sampling sites were selected exclusively from the coastal areas of Guangdong Province, all experiencing the same subtropical monsoon climate(Ding et al., 2022). This study aims to: (1) analyze microbiota across diverse ancestral hall surfaces and assess their potential functions; (2) compare microbial diversity between damaged and undamaged surfaces to evaluate the protective effects of paint; (3) characterize microbial communities and their correlation with environmental factors; (4) validate amplicon sequencing results through strain isolation experiments; (5) assess the antimicrobial sensitivity of microorganisms to four biocides. These findings will deepen our understanding of ancestral hall microbial weathering and provide valuable insights for safeguarding wooden structures in Guangdong Province or similar subtropical monsoon climate conditions.

## 2. Materials and methods

### 2.1. Sample sites

Sampling was carried out at nine wooden ancestral halls in Guangdong Province, located in the southeast coastal region of China (Figure 1). These ancestral halls share a typical subtropical monsoon climate but vary in city location, enabling observation of microbiome commonalities across different areas within the same climatic conditions. Based on distinct visitor flow rates, the halls were categorized into three groups: the urban group (Urb_CHEN, Urb_GUAN, Urb_SAN) situated in the central city of Guangzhou, with an average hourly population of 10,000 people; the suburban group (Sub_LIN, Sub_DONG, Sub_LIE) located on the outskirts of Guangzhou, with an average hourly population of 100 people; and the rural district group (RD_QING, RD_FU, RD_MING) situated in the country areas of Jieyang, with an average hourly population of less than 10.

**Figure 1.**
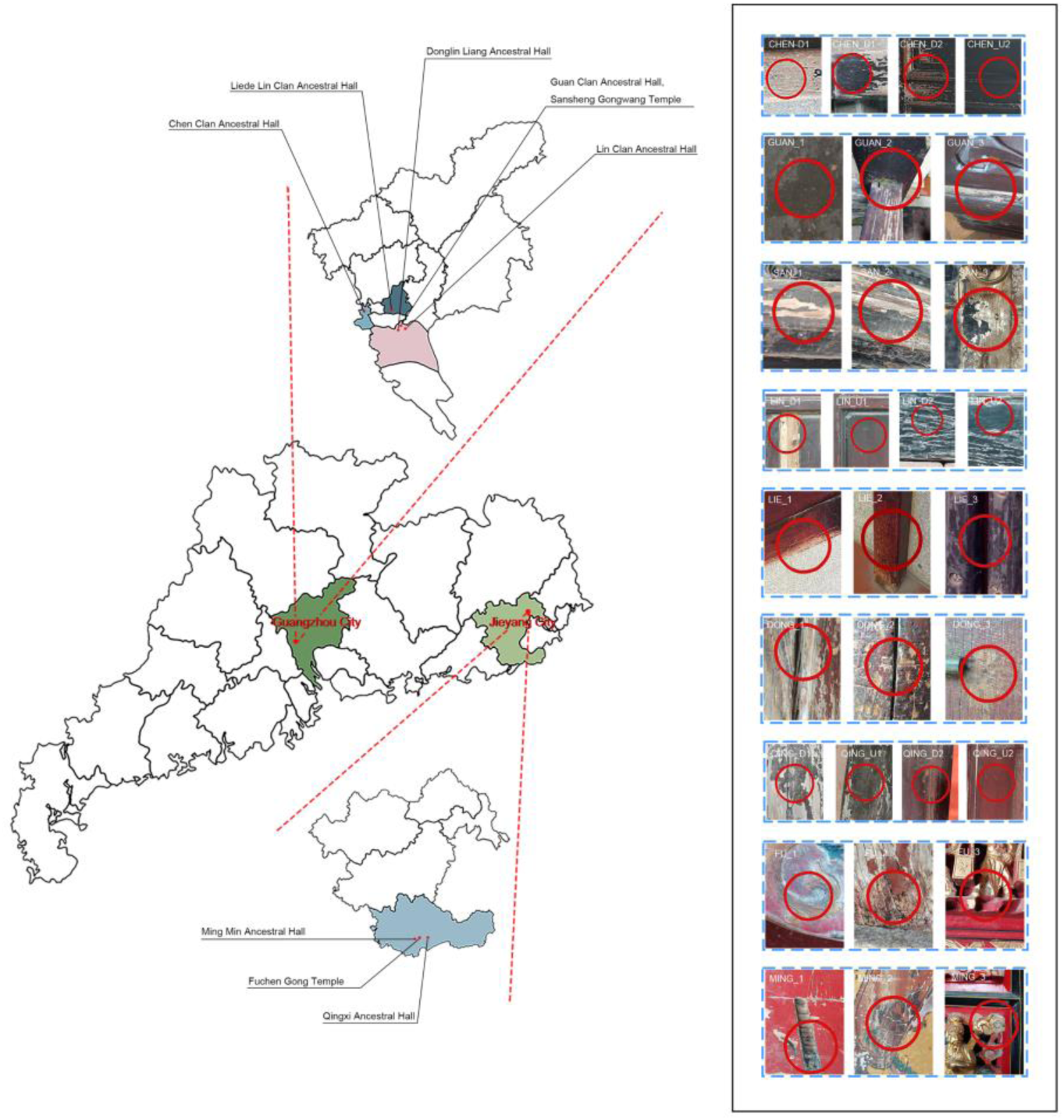
Characteristics of the samples. The location of nine ancestral halls and their corresponding sampling sites are provided. Samples CHEN_D1, CHEN_U1, LIN_D2, LIN_U2, and FU_2 were collected from wooden thresholds. Samples CHEN_D2, CHEN_U2, GUAN_1, SAN_3, LIN_D1, LIN_U1, DONG_3, LIE_1, and MING_2 were sampled from wooden doors. Sample GUAN_2 was taken from a wooden railing. Samples GUAN_3, QING_D2, and QING_U2 were obtained from wooden folding screens. Samples SAN_1 and SAN_2 were taken from wooden beams. Sample LIE_2 was sampled from a wooden table corner. Sample LIE_3 was obtained from a wooden bolt. Samples DONG_1, DONG_2, QING_D1, and QING_U1 were sampled from wooden pillars. Sample FU_1 was obtained from a wooden incense burner. Samples FU_3 and MING_3 were sampled from shrines. Sample MING_1 was obtained from a sacrificial table.

We selected two ancestral halls from each group and took three samplings from the damaged external surfaces of Urb_GUAN (GUAN_1, GUAN_2, GUAN_3), Urb_SAN (SAN_1, SAN_2, SAN_3), Sub_DONG (DONG_1, DONG_2, DONG_3), Sub_LIE (LIE_1, LIE_2, LIE_3), RD_FU (FU_1, FU_2, FU_3), and RD_MING (MING_1, MING_2, MING_3). Additionally, one ancestral hall from each group provided two pairs of samplings from damaged and undamaged external surfaces: Urb_CHEN (CHEN_D1, CHEN_U1, CHEN_D2, CHEN_U2), Sub_LIN (LIN_D1, LIN_U1, LIN_D2, LIN_U2), and RD_QING (QING_D1, QING_U1, QING_D2, QING_U2). The damaged sites are adjacent to their undamaged counterparts with the same environmental conditions, both being sampled to achieve a direct comparison of microbiome colonization.

### 2.2. Environmental data recording

Sampling was conducted during summer because microbial metabolism and reproduction are more active at this time due to its characteristic high temperatures and heavy rainfall; while winters are milder with less precipitation. Choosing a random summer day serves to represent the overall environmental conditions throughout the entire summer season, given the relatively stable environmental conditions within each season, which is also a typical character of subtropical monsoon climate. Moreover, Ancestral halls offer a semi-open, semi-indoor environment, exposed yet shielded by eaves and walls, ensuring relatively stable conditions for surface microbial growth.

Measurements of relative humidity (RH), temperature (C), and light (lux) were recorded at each site using SMART SENSOR AS817 Humidity & Temperature Meter and AS803 Digital Lux Meter, respectively. Wood moisture content was assessed with SMART SENSOR AS981 Moisture Meter. Additionally, hourly visitor counts were collected at the entrance of the ancestral hall and obtained from staff records.

Details about the sampling location, including relative altitude from ground level and wood face direction (aspect), were noted alongside the aforementioned environmental factors. Data and photographs were collected to enable precise future resampling if needed.

### 2.3. Surface sampling

Sampling was performed using sterile swabs. To obtain samples a sterile swab was dipped in sterile M9 salts and wiped over a 5 cm square region of the surface. Each sample was allocated an individual tube of fresh M9 salts to prevent cross-contamination between sampling sites. At all stages of sampling, nitrile gloves were worn to prevent skin microbiota contamination. Samples were refrigerated at 4°C for further culturing or sequencing.

### 2.4. Amplicon Sequencing Analysis

The 16S rRNA V3–V4 amplicon was amplified using 2×Hieff® Robust PCR Master Mix (Yeasen,10105ES03, China) and two universal bacterial 16S rRNA gene amplicon PCR primers: Nobar_341F (5’-CCTACGGGNGGCWGCAG-3’) and Nobar_805R (5’-GACTACHVGGGTATCTAATCC -3’). For the reaction setup, microbial DNA (10ng/µl) of 2µl, amplicon PCR forward primer (10µM) of 0.5µl, amplicon PCR reverse primer (10 µM) of 0.5µl, 2×Taq Master Mix (Sangon Biotech, B639295, China) of 12.5µl, and 9.5µL double distilled water (ddH2O) were combined to a total volume of 25µl. PCR was carried out in a T100 Thermal Cycler (1861096, USA) with the following program: initial denaturation at 95 °C for 3 min, followed by 5 cycles of denaturation at 95 °C for 30 s, annealing at 55 °C for 30 s, elongation at 72 °C for 1min 30 s, then 35 cycles of denaturation at 95 °C for 30 s, annealing at 55 °C for 30 s, elongation at 72 °C for 1min 30 s, and a final extension at 72 °C for 10 min.

The 18S rRNA V4 amplicon was amplified using 2×Hieff® Robust PCR Master Mix (Yeasen, 10105ES03, China) and universal fungal 18S rRNA V4 gene amplicon PCR primers: 18SV4-forward primer (5’-GGCAAGTCTGGTGCCAG-3’) and 18SV4-reverse primer (5’-ACGGTATCTRATCRTCTTCG-3’). For the reaction setup, microbial DNA (10ng/µl) of 2µl, amplicon PCR forward primer (10µM) of 1µl, amplicon PCR reverse primer (10 µM) of 1µl, and 2×Hieff® Robust PCR Master Mix of 15µl were combined to a total volume of 30µl. PCR was performed in an Applied Biosystems 9700 thermal instrument (USA) with the following program: initial denaturation at 94 °C for 3 min, followed by 5 cycles of denaturation at 94 °C for 30 s, annealing at 45 °C for 20 s, elongation at 65 °C for 30 s, then 20 cycles of denaturation at 94°C for 20 s, annealing at 55 °C for 20 s, elongation at 72 °C for 30 s, and a final extension at 72 °C for 5 min.

Samples were sent to Sangon BioTech (Shanghai) for library construction, employing universal Illumina adaptors and indexes. Prior to sequencing, DNA concentration of each PCR product was quantified using Qubit® 4.0 Green double-stranded DNA assay and assessed for quality with a bioanalyzer (Agilent 2100, USA). Libraries, based on coverage needs, were pooled for sequencing in one run, with amplicons combined in equimolar ratios according to their concentrations. Sequencing was conducted using Illumina MiSeq system (Illumina MiSeq, USA) per manufacturer’s guidelines. Sequence processing, operational taxonomic unit (OTU) clustering, Representative tags alignment, and biological classification were performed. PEAR software (version 0.9.8) assembled the paired short Illumina readings from overlap, generating fasta and qual files for standard analysis. Usearch software (version 11.0.667) clustered effective tags into OTUs with ≥97% similarity, filtering out chimeric sequences and singleton OTUs (single-read OTUs). Remaining sequences were sorted into respective samples based on OTUs, selecting the most abundant tag sequence per cluster. Taxonomic classification of bacterial and fungal OTU representative sequences was conducted using RDP Database and UNITE fungal ITS Database through sequence blasting.

### 2.5. Bioinformatics analysis

The alpha diversity indices, including Chao 1, Simpson, and Shannon indices, were quantified with the R package Vegan to demonstrate the diversity of microbial community.

### 2.6. Function prediction

The bacteria’s functional prediction analysis was performed using PICRUSt (v1.1.4) software, aligning sequenced 16S rRNA gene data with a reference genome database. This enabled the inference of potential gene families associated with each OTU. By integrating gene family data and relative abundances of microbial OTUs, PICRUSt predicted gene family abundances in our samples, offering insights into functional potential based on 16S rRNA gene sequences.

### 2.7. Isolation, and identification of microorganisms

Microorganisms on the swabs were separated from the swabs into the solution by vortexing at 180rpm at 30°C for 3 hours. The resulting suspension was plated on non-selective media—BPM (Beef extract Peptone Medium) plates for bacteria and PDA (Potato Dextrose Agar Medium) plates (with added 0.1g/L chloramphenicol) for fungi—and cultured for 2-7 days at 28°C . Colonies were separated based on color and morphology, and the isolating process was repeated to obtain purebred strains. These strains underwent sequencing for genus and species identification.

The extraction of genomic DNA from isolated strains was conducted using the CTAB Extraction Method. (Moller et al., 1992).

The 16S rRNA V3–V4 amplicon was amplified using 2×Taq Master Mix (Sangon Biotech, B639295, China) and two universal bacterial 16S rRNA gene amplicon PCR primers: 27F (5′-AGAGTTTGATCCTGGCTCAG-3′) and 1492R (5′-GGTTACCTTGTTACGACTT-3′). The reaction setup included 2µL of microbial DNA (10ng/µl), 0.5µL of amplicon PCR forward primer (10µM), 0.5µL of amplicon PCR reverse primer (10 µM), and 12.5µL of 2×Taq Master Mix, with a total volume of 25µL including 9.5µL of ddH2O. PCR was carried out in a T100 Thermal Cycler (1861096, USA) with the following program: initial denaturation at 95 °C for 3 min, followed by 35 cycles of denaturation at 95 °C for 30 s, annealing at 55 °C for 30 s, elongation at 72 °C for 1 min 30 s, and a final extension at 72 °C for 10 min.

The ITS amplicon was amplified using 2×Taq Master Mix (Sangon Biotech, B639295, China) with two universal ITS amplicon PCR primers: ITS1-forward primer (5’-TCCGTAGGTGAACCTGCGG-3’) and ITS4-reverse primer (5’-TCCTCCGCTTATTGATATGC-3’). For the reaction setup, 2µL of microbial DNA (10ng/µL), 0.5µL of amplicon PCR forward primer (10µM), 0.5µL of amplicon PCR reverse primer (10 µM), and 12.5µL of 2×Taq Master Mix were combined, with a total volume of 25µL including 9.5µL of ddH2O. PCR was conducted in a T100 Thermal Cycler (1861096, USA) following this program: initial denaturation at 95 °C for 3 min, followed by 35 cycles of denaturation at 95 °C for 30 s, annealing at 55 °C for 30 s, elongation at 72 °C for 1 min, and a final extension at 72 °C for 10 min.

### 2.8. Identification of cellulolytic microbes and ligninolytic microbes

odium carboxymethyl cellulose (CMC-Na) and guaiacol were employed to screen for cellulolytic and ligninolytic microbes, respectively. The composition of the media used for screening is detailed as follows: CMC-degrading agar medium for bacteria (per liter): CMC-Na 15.0g, Na2HPO4·7H2O 12.8g, KH2PO4 3.0g, NaCl 0.5g, NH4Cl 1.0g, agar 15.0g; CMC-degrading agar medium for fungi (per liter): CMC-Na 2.0g, KH2PO4 1.0g, MgSO4 0.5g, (NH4)2SO4 2.0g, NaCl 0.5g, agar 18.0g; lignin-degrading agar medium for bacteria (per liter): 9mM guaiacol, yeast extract 50mg, Na2HPO4·7H2O 12.8g, KH2PO4 3.0g, NaCl 0.5g, NH4Cl 1.0g, agar15.0g; lignin-degrading agar medium for fungi (per liter): 9mM liquid guaiacol, MgSO4 0.5g, KH2PO4 1.0g, Na2HPO4 0.2g.

CMC-Na was the sole carbon source in the CMC-degrading medium to ensure cellulose degradation as the only carbon source for microbial growth. Additionally, yeast extract was added in small quantities to the lignin-degrading medium as an auxiliary carbon source. Plates were incubated at 28°C for 48 hours (bacteria) or 96 hours (fungi), and the diameter of the degradation circles was measured in millimeters using a scale ruler.

### 2.9. The disk-diffusion test of biocides

The study evaluated microbial sensitivity to Isothiazolinone, NaClO, Thymus Vulgaris Essential Oil, and Cinnamon Essential Oil using the disk diffusion method. Microbes were cultivated on BPM agar for bacteria (48 hours at 28°C) and PDA agar for fungi (96 hours at 28°C). Sterile paper disks of 6-mm diameter were soaked in respective biocides for an hour and then placed on the culture mediums. The antimicrobial effect was determined by measuring the diameter of inhibition zones using a scale ruler, indicating the extent of antimicrobial activity for each concentration.

Two forms of microorganisms were assessed: isolated strains and microbial assembly. Both forms underwent testing with four biocides at a concentration of 1% to ascertain their fungicidal or bactericidal activity. The biocides displaying the most potent antimicrobial effect were further investigated through a concentration gradient experiment. Seven concentrations (1%, 0.5%, 0.25%, 0.125%, 0.1%, 0.05%, 0.01%) were prepared, employing the disk diffusion method to individually assess the antimicrobial effects on the microbial community.

## 3. Results

### 3.1. Characteristics of the samples

Nine ancestral halls with varying visitor flow rates were sampled, and environmental factors at these sites were measured (Table 1, Table S1, and Figure 1). Despite being typically protected by paint, weathering has damaged some surfaces, leading to flaked paint, exposed wood, and visible biocolonization—defined here as ’damaged sites.’ Observations include dark green residue on the wooden railing (GUAN_2), white biofouling on surfaces like the wooden door (SAN_3), wooden pillar (DONG_1), and shrine (FU_3), as well as notable black spots on the threshold (FU_2) and sacrificial table (MING_1), and yellow patches on the wooden screen (QING_2).

**Table 1:**
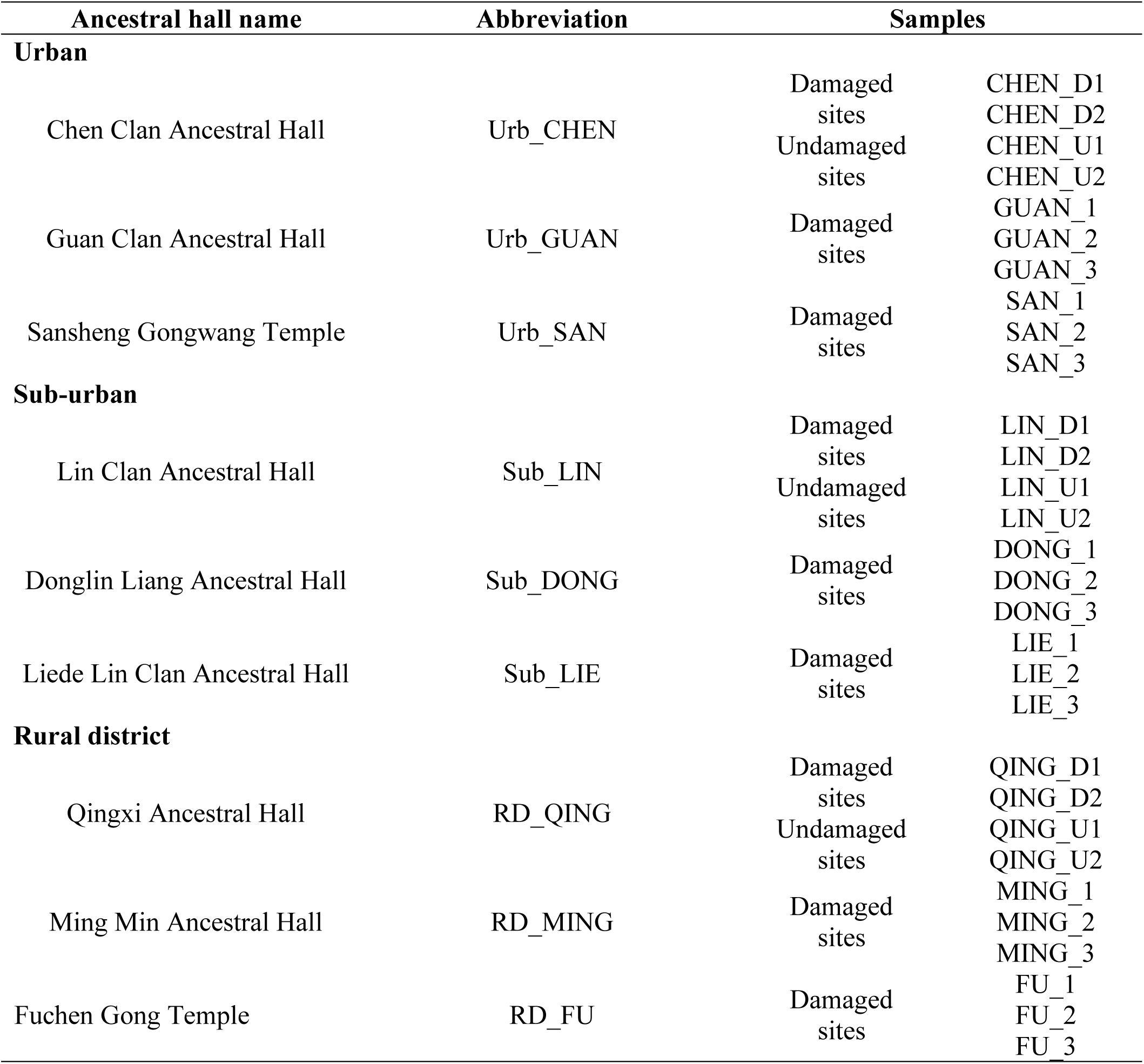
Abbreviation of the sampling ancestral halls and their samples.

| Ancestral hall name | Abbreviation | Samples |  |
| --- | --- | --- | --- |
| Urban |  |  |  |
| Chen Clan Ancestral Hall | Urb_CHEN | Damaged sites | CHEN_D1 |
|  |  |  | CHEN_D2 |
|  |  | Undamaged sites | CHEN_U1 |
|  |  |  | CHEN_U2 |
| Guan Clan Ancestral Hall | Urb_GUAN | Damaged sites | GUAN_1 |
|  |  |  | GUAN_2 |
|  |  |  | GUAN_3 |
| Sansheng Gongwang Temple | Urb_SAN | Damaged sites | SAN_1 |
|  |  |  | SAN_2 |
|  |  |  | SAN_3 |
| Sub-urban |  |  |  |
| Lin Clan Ancestral Hall | Sub_LIN | Damaged sites | LIN_D1 |
|  |  |  | LIN_D2 |
|  |  | Undamaged sites | LIN_U1 |
|  |  |  | LIN_U2 |
| Donglin Liang Ancestral Hall | Sub_DONG | Damaged sites | DONG_1 |
|  |  |  | DONG_2 |
|  |  |  | DONG_3 |
| Liede Lin Clan Ancestral Hall | Sub_LIE | Damaged sites | LIE_1 |
|  |  |  | LIE_2 |
|  |  |  | LIE_3 |
| Rural district |  |  |  |
| Qingxi Ancestral Hall | RD_QING | Damaged sites | QING_D1 |
|  |  |  | QING_D2 |
|  |  | Undamaged sites | QING_U1 |
|  |  |  | QING_U2 |
| Ming Min Ancestral Hall | RD_MING | Damaged sites | MING_1 |
|  |  |  | MING_2 |
|  |  |  | MING_3 |
| Fuchen Gong Temple | RD_FU | Damaged sites | FU_1 |
|  |  |  | FU_2 |
|  |  |  | FU_3 |

### 3.2. Taxonomic Analysis of the diversity of microbial community

The taxonomic analysis revealed a total of 1467 OTUs from the 16S sequences and 367 from the 18S sequences after quality filtering. Despite three PCR repetitions on the MING_3 sample, the DNA concentration remained insufficient for library construction of amplicons. Alpha diversity metrics for both prokaryotic and eukaryotic communities—Shannon, Chao, Ace, Simpson, and Shannon evenness indices—were calculated to assess sample biodiversity. This analysis indicates limited fungal species and varying bacterial abundance. The coverage index, part of alpha diversity, evaluates observed species against the total pool. Our study demonstrates a coverage index close to 1, signifying a comprehensive representation of species in our sample (Table S2 and Table S3).

We visualized the predominant prokaryotic and eukaryotic organisms in samples from damaged surfaces, revealing commonalities across all ancestral halls. At the phylum level, *Firmicutes* and *Proteobacteria*, the two dominant prokaryotic phyla, collectively represent over 84% of the total OTU relative abundance in all nine ancestral halls (Figure 2A, Figure S1A, and Table S4).

**Figure 2.**
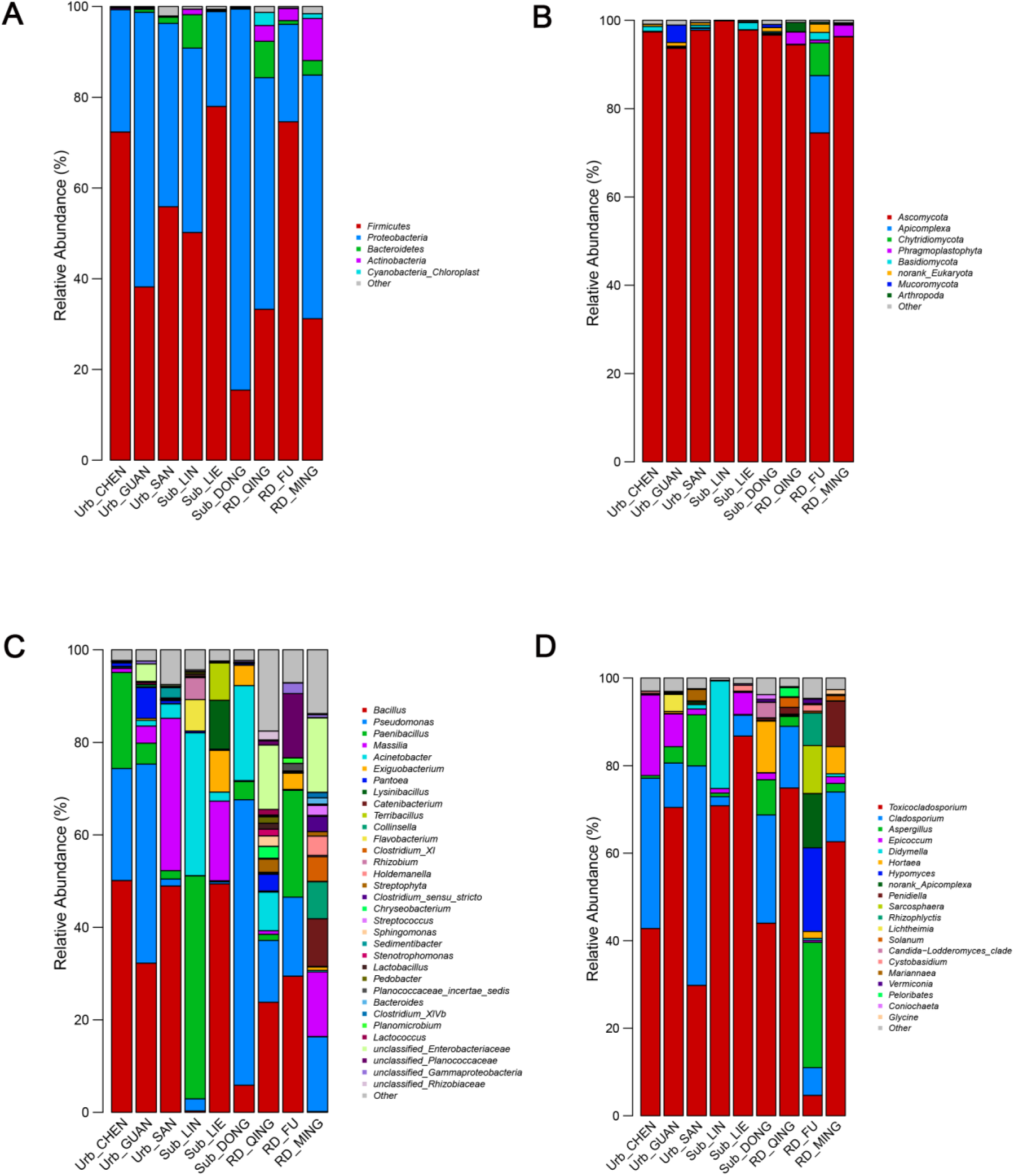
Determining the diversity of microbial community. (A, B, C and D) Percentage of amplicon reads mapped to each ancestral hall. Phylum level taxonomy and genus level taxonomy of microbiome on the damaged surfaces of nine ancestral halls under the same climate conditions, based on 16S amplicon sequencing and 18S amplicon sequencing. (A) Bacterial phyla. (B) Fungal phyla. (C) Bacterial genera. (D) Fungal genera.

*Ascomycota* emerges as the dominant eukaryotic phylum, accounting for more than 90% abundance in eight of the nine ancestral halls (excluding RD_FU, which stands at 74.53%) (Figure 2B, Figure S1B, and Table S5). Zooming in to the genus level, four dominant bacterial genera—*Bacillus* (except in Sub_LIN and RD_MING, where abundance is below 1%), *Pseudomonas* (except in Sub_LIE, abundance below 1%), *Paenibacillus* (except in Sub_LIE and RD_MING, abundance below 1%), and *Acinetobacter* (except in Urb_CHEN, RD_FU, and RD_MING, abundance below 1%)—were identified (Figure 2C, Figure S1C, and Table S6). Among eukaryotic genera, *Toxicocladosporium*, *Cladosporium*, *Aspergillus* (except in Urb_CHEN, Sub_LIN, and Sub_LIE, abundance below 1%), and *Epicoccum* (RD_QING and RD_FU, abundance below 1%) demonstrated significant abundance across all nine ancestral halls (Figure 2D, Figure S1D, and Table S7).

### 3.3. Comparison of microbial abundance between damaged sites and undamaged sites

There are conflicts in discussion over whether species are associated with damaged or undamaged surfaces and whether they play an important role in biodeterioration(Skipper et al., 2022). Therefore, to investigate into the role of microorganisms in biodeterioration, we took pairs of closely related damaged and undamaged samples exposed to identical climatic conditions.

we selected three ancestral halls—the Chen Clan Ancestral Hall, the Lin Clan Ancestral Hall, and the Qingxi Ancestral Hall—and collected two pairs of samples from damaged and undamaged surfaces at each location. These damaged sites are adjacent to their undamaged counterparts, sharing identical environmental conditions. This sampling strategy allows for a direct comparison of microbiome colonization. Heatmaps were utilized to highlight differences in microbial taxa abundances between these groups.

Among prokaryotic microorganisms, nine phyla (Figure S2A) and 38 bacterial genera (Figure S2C) were present in both damaged and undamaged samples. Four phyla—*Actinobacteria, Bacteroidetes, Firmicutes,* and *Proteobacteria*—along with four bacterial genera—*Bacillus, Paenibacillus, Pseudomonas,* and *Pantoea*—showed notably high abundance across all samples (Figure S3, Figure S1E, and Figure S4). *Firmicutes, Paenibacillus, Bacillus,* and *Acinetobacter* were more abundant in damaged samples, while *Proteobacteria* and *Actinobacteria* were more prevalent in undamaged samples. Regarding eukaryotic microorganisms, seven phyla (Figure S2B) and 13 fungal genera (Figure S2D) were shared among all damaged and undamaged samples. Five phyla—*Chlorophyta, Ascomycota, Basidiomycota, Phragmoplastophyta,* and *Ascomycota*—along with four fungal genera—*Toxicocladosporium, Epicoccum, Cladosporium,* and *Aspergillus*—showed differential abundance across all samples, with *Toxicocladosporium* exhibiting the highest abundance (Figure S1F, Figure S5, and Figure S6). *Ascomycota* was more prevalent in damaged samples, while *Phragmoplastophyta* was more abundant in undamaged samples. Notably, *Cladosporium* exhibited significant higher abundance in damaged samples (except for the Lin Clan Ancestral Hall).

### 3.4. The relationship of the abundance of microbial community and environmental factors

To assess the impact of environmental factors—such as relative humidity, temperature, light, wood moisture content, and visitor flow rate—on microbial abundance and diversity, a Spearman correlation heatmap was employed. Taking visitor flow rate as an example, correlations were observed between microbial abundance and this factor. For prokaryotes, *Sedimentibacter, Trichococcus, Terribacillus, Pantoea,* and *Proteiniphilum* showed positive correlations with visitor flow rate, while *Streptophyta, Lactococcus, Sphingomonas,* and *Flavobacterium* exhibited negative correlations (Figure S7). Among eukaryotes, *Rhizopus* and *Lichtheimia* displayed positive correlations with visitor flow rate, whereas *Glycine* and *Solanum* exhibited negative correlations (Figure S8).

Beyond the visitor flow rate, other environmental factors also contributed to the composition of microbial community. For instance, relative humidity showed significant correlations with the abundance of various fungi. Additionally, temperature and wood moisture content impacted the abundance of certain bacteria. The abundance and diversity of the microbial community is the consequence of collaborative interaction among all environmental factors.

### 3.5. Genetic function prediction for the microbial community

Through KEGG analysis using PICRUSt, we inferred a potential integrated cellulose decomposition pathway across all sample sites, illustrating a hypothetical breakdown of cellulose into cellulose dextrin via metabolic pathways K01179 and K01188, succeeded by further decomposition into cellobiose (Figure 3A).

**Figure 3.**
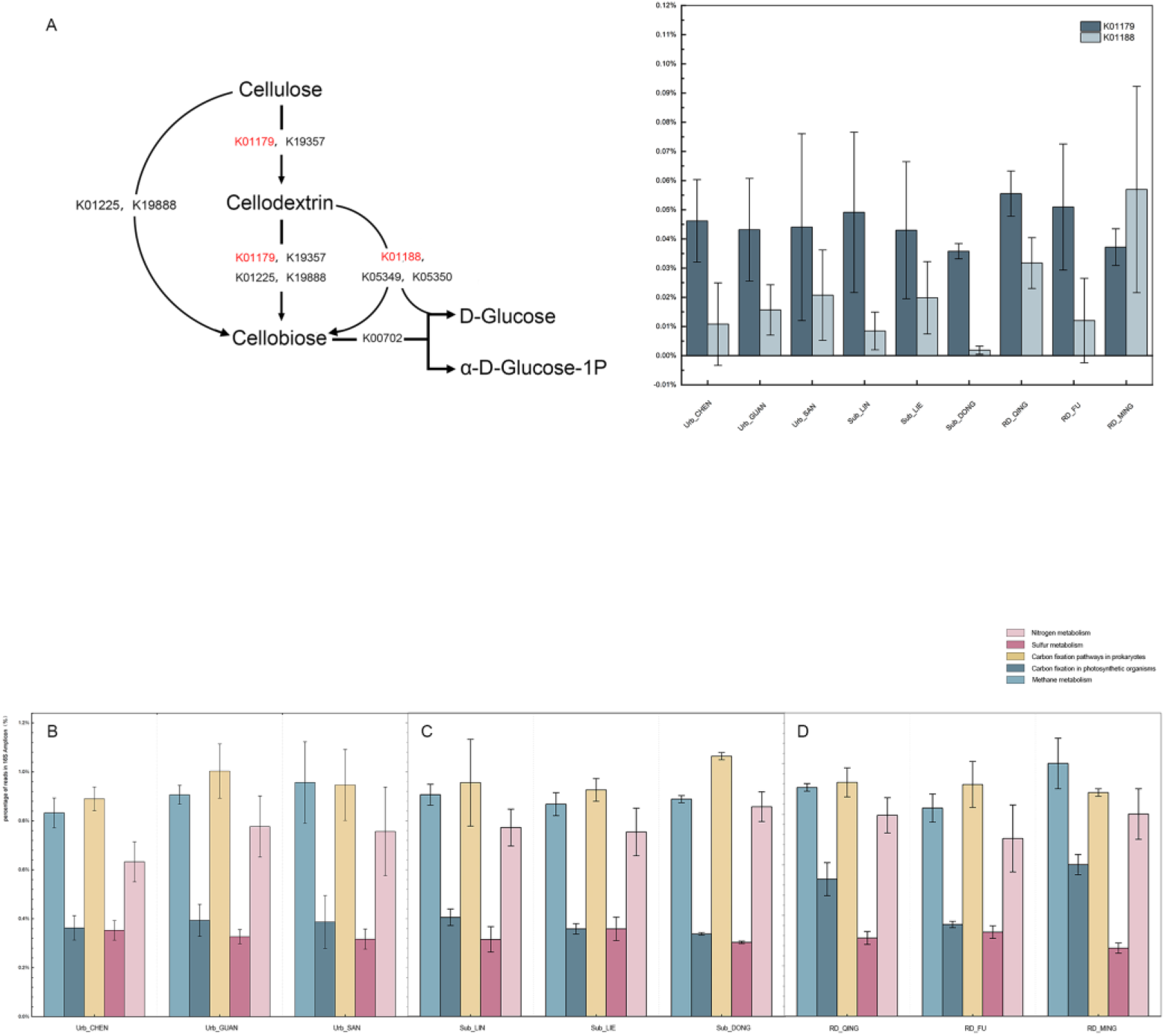
Genetic function prediction for the microbial community. (A) The percentage of the cellulose degradation pathways inferred at the microbiome on the surfaces of ancestral halls based on KEGG Orthology, some of which that marked in red were detected in our study. (B, C and D) The major functions driving geomicrobiological energy metabolism of microbiome communities on urban (A), sub-urban (B), and rural (C) ancestral halls. Totally five functions are summarized through KEGG Pathway. A total of five functions are included, and all abundances are reported as percentages of amplicon reads. Vertical lines indicate the standard deviation.

Additionally, we summarized the relative abundance of five microbial biological cycle functions associated with fundamental nutrient cycling. KEGG annotation revealed relatively high abundance of prokaryotic carbon fixation and nitrogen metabolism in all samples. Moreover, simultaneous sulfur metabolism, photosynthetic carbon fixation, and methane metabolism were evident in this microbiota, indicating an active metabolic process within the bacterial community in the samples (Figure 3B, Figure 3C, and Figure 3D).

### 3.6. Verification of the degrading ability of microorganisms from Urb_CHEN based on conventional culturing methods

The amplicon sequencing revealed a highly similar microbial composition across all ancestral halls. Consequently, one ancestral hall, Urb_CHEN, was chosen for further microorganism isolation. Using conventional culturing methods, 8 bacterial species and 7 fungal species were isolated from the Urb_CHEN samples (Table 2). All isolates underwent testing for cellulose and lignin degradation ability, except for the *Staphylococcus* strain due to its pathogenicity, which precluded further experimentation.

**Table 2:** Molecular identification of strains isolated from the four samples of Urb_CHEN.

| Nucleotide Blast reference strains |  |  |  |  |
| --- | --- | --- | --- | --- |
|  | Clost relative strain | Accession number | Phylum | Similarity (%) |
| **Bacteria** |  |  |  |  |
| CHEN_B1 | *Pseudomonas* sp. | MN043751.1 | Proteobacteria | 99.79% |
| CHEN_B2 | *Pantoea dispersa* strain | MT921704.1 | Proteobacteria | 99.86% |
| CHEN_B3 | *Paenibacillus* sp. | JF768727.1 | firmicutes | 99.72% |
| CHEN_B4 | *Bacillus cereus* strain | MN691548.1 | Firmicutes | 98.21% |
| CHEN_B5 | *Priestia megaterium* strain | MF431767.1 | Firmicutes | 100.00% |
| CHEN_B6 | *Staphylococcus hominis* strain | KY992547.1 | Firmicutes | 100.00% |
| CHEN_B7 | *Curtobacterium* sp. | MK704290.1 | Actinobacteria | 99.72% |
| CHEN_B8 | *Microbacterium oleivorans* strain | JQ660049.1 | Actinobacteria | 99.65% |
| **Fungi** |  |  |  |  |
| CHEN_F1 | *Rhodotorula mucilaginosa* strain | ON954707.1 | Basidiomycota | 99.84% |
| CHEN_F2 | *Cystobasidium* sp. | AF444619.1 | Basidiomycota | 99.32% |
| CHEN_F3 | *Aureobasidium pullulans* strain | KT352844.1 | Ascomycota | 98.23% |
| CHEN_F4 | *Cladosporium* sp. | OK242732.1 | Ascomycota | 99.82% |
| CHEN_F5 | *Trichoderma* sp. | ON248245.1 | Ascomycota | 99.84% |
| CHEN_F6 | *Daldinia* sp. | MK311339.1 | Ascomycota | 99.83% |
| CHEN_F7 | *Aspergillus sydowii* strain | OP237096.1 | Ascomycota | 99.65% |

<sup>a</sup>“CHEN\_B1 ~ CHEN\_B8” and “CHEN\_F1 ~ CHEN\_F7” indicate strains were identified by 16S and ITS rDNA gene. CHEN\_B2, CHEN\_B4, CHEN\_B5, CHEN\_B6, CHEN\_B8, CHEN\_F1, CHEN\_F3, and CHEN\_F7 were successfully annotated to the species level, while the remaining strains could only be annotated to the genus level.

Among all the bacteria, *Pseudomonas* sp., *Pantoea dispersa*, *Paenibacillus* sp., *Bacillus cereus*, and *Priestia megaterium* demonstrated both cellulose and lignin degradation abilities (Figure 4A-E). *Curtobacterium* sp. showed lignin degradation ability but not cellulose degradation (Figure 4F). *Microbacterium oleivorans* exhibited no degradation capability for either lignin or cellulose. Among the fungi, *Aureobasidium pullulans*, *Cladosporium* sp., *Trichoderma* sp., *Daldinia* sp., and *Aspergillus sydowii* could degrade cellulose but not lignin, evident from distinct hydrolytic zones on plates (Table 3 and Figure S9A-E). Conversely, *Rhodotorula mucilaginosa* and *Cystobasidium* sp. displayed no capability for degrading either lignin or cellulose.

**Figure 4.**
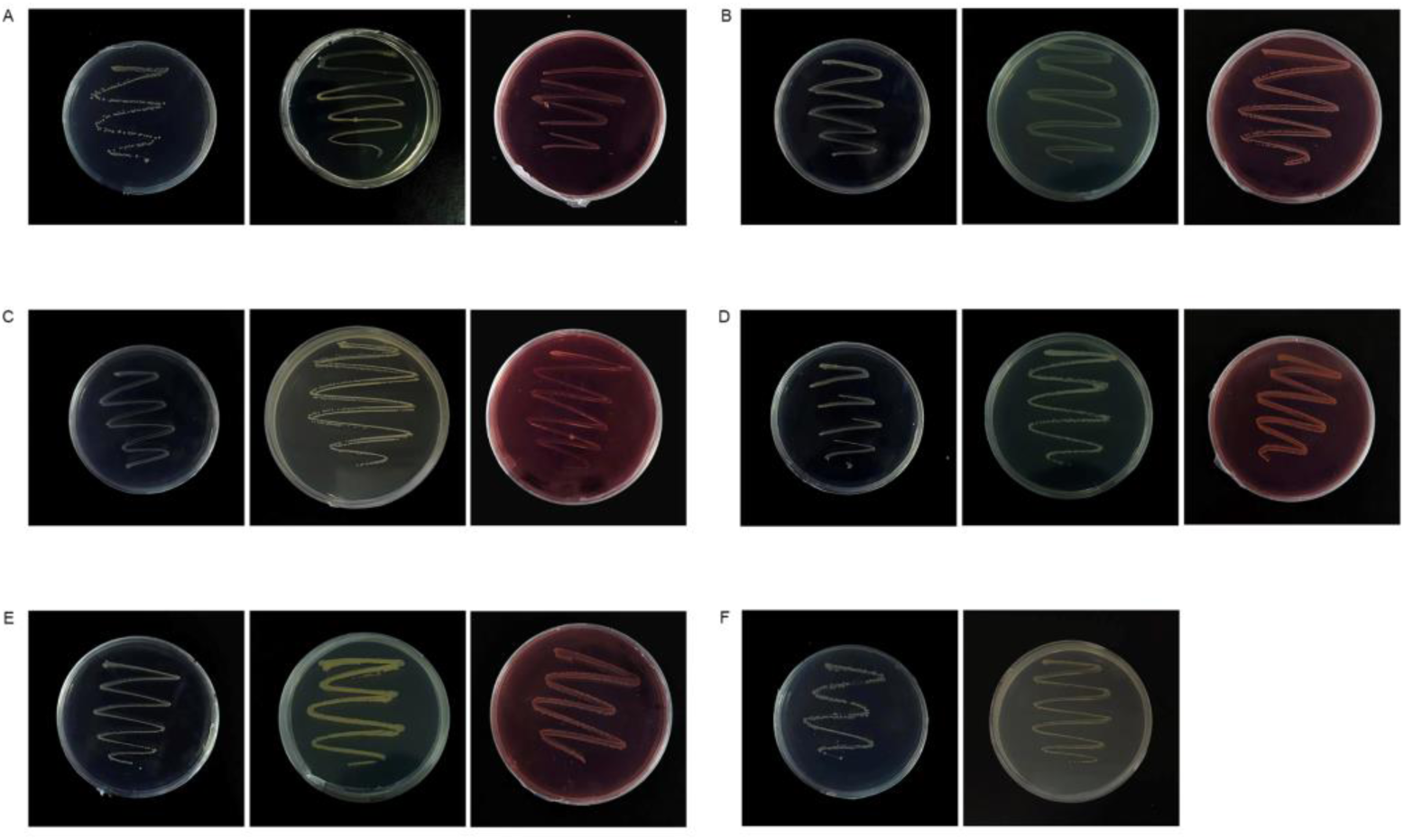
Verification of the degrading ability of microorganisms from Urb_CHEN based on conventional culturing methods. The three media for each isolate, from left to right, are the lignin-degrading agar medium, the LB medium, and the CMC-degrading agar medium. Bacterial isolates capable of utilizing guaiacol on the lignin-degrading agar medium and cellulose on the CMC-degrading agar medium for two days are shown. (A)*Pseudomonas* sp. (B) *Pantoea dispersa* strain. (C) *Paenibacillus* sp. (D) *Bacillus cereus* strain. (E) *Priestia megaterium* strain. Bacterial isolates that can only utilize guaiacol on the lignin-degrading agar medium for two days. (F) *Curtobacterium* sp. The *Microbacterium oleivorans* strain which cannot grow on lignin-degrading agar medium or CMC-degrading agar medium does not be shown.

**Table 3:** Hydrolytic zones of the fungal isolates on CMC-degrading agar medium after three days of incubation.

| Fungi | Hydrophytic zones diameter (mm) |
| --- | --- |
| CHEN_F1 | No degradation |
| CHEN_F2 | No degradation |
| CHEN_F3 | 5.00±0.00 |
| CHEN_F4 | 8.08±1.18 |
| CHEN_F5 | 23.67±3.51 |
| CHEN_F6 | 9.67±0.76 |
| CHEN_F7 | 5.33±0.58 |

872 <sup>a</sup>Data are represented as mean ± SEM.

### 3.7. Antimicrobial sensitivity test of the microorganisms from Urb_CHEN

Using biocides is a common method to prevent biodeterioration on the surface of cultural relics. Four biocides—Isothiazolinone, Sodium hypochlorite, Thymus Vulgaris Essential Oil, and Cinnamon Essential Oil—were selected. Antimicrobial sensitivity tests were conducted on microorganisms obtained from Urb_CHEN samples using the agar disk-diffusion method. Two forms of microorganisms were tested: isolated and microbial assembly forms. As we mentioned in the Materials and Methods section, Samples were collected by swabbing ancestral hall surfaces with a sterile swab dipped in M9 salts solution, followed by vortexing to separate microorganisms into a suspension. This suspension was plated onto media for growth, forming the ’microbial assembly’ referenced in our study.

In biocidal testing on isolated strains, Isothiazolinone at a 1% concentration showed notable antibacterial effects against all 8 bacterial isolates and all 7 fungi isolates. In contrast, Sodium hypochlorite at the same concentration exhibited inhibitory effects on specific fungi isolates only (Figure 5A and B, Table 4). In testing on microbial assembly, Isothiazolinone exhibited complete antimicrobial effects on fungi community and notably inhibited bacteria community, as evidenced by significant inhibition zones (25.50±0.50mm, 29.00±1.73mm, 28.17±1.76mm, and 48.33±2.08mm for samples CHEN_D1, CHEN_U1, CHEN_D2, and CHEN_U2). Sodium hypochlorite showed limited antibacterial function but displayed inhibitory effects on fungi community, as indicated by inhibition zone diameters (26.67±3.06mm, 24.00±3.46mm, 24.33±2.52mm, and 19.33±1.15mm for samples CHEN_D1, CHEN_U1, CHEN_D2, and CHEN_U2). However, Thymus Vulgaris Essential Oil and Cinnamon Essential Oil did not significantly suppress microbial growth at a 1% (v/v) concentration for both isolates and microbial assembly.

**Figure 5.**
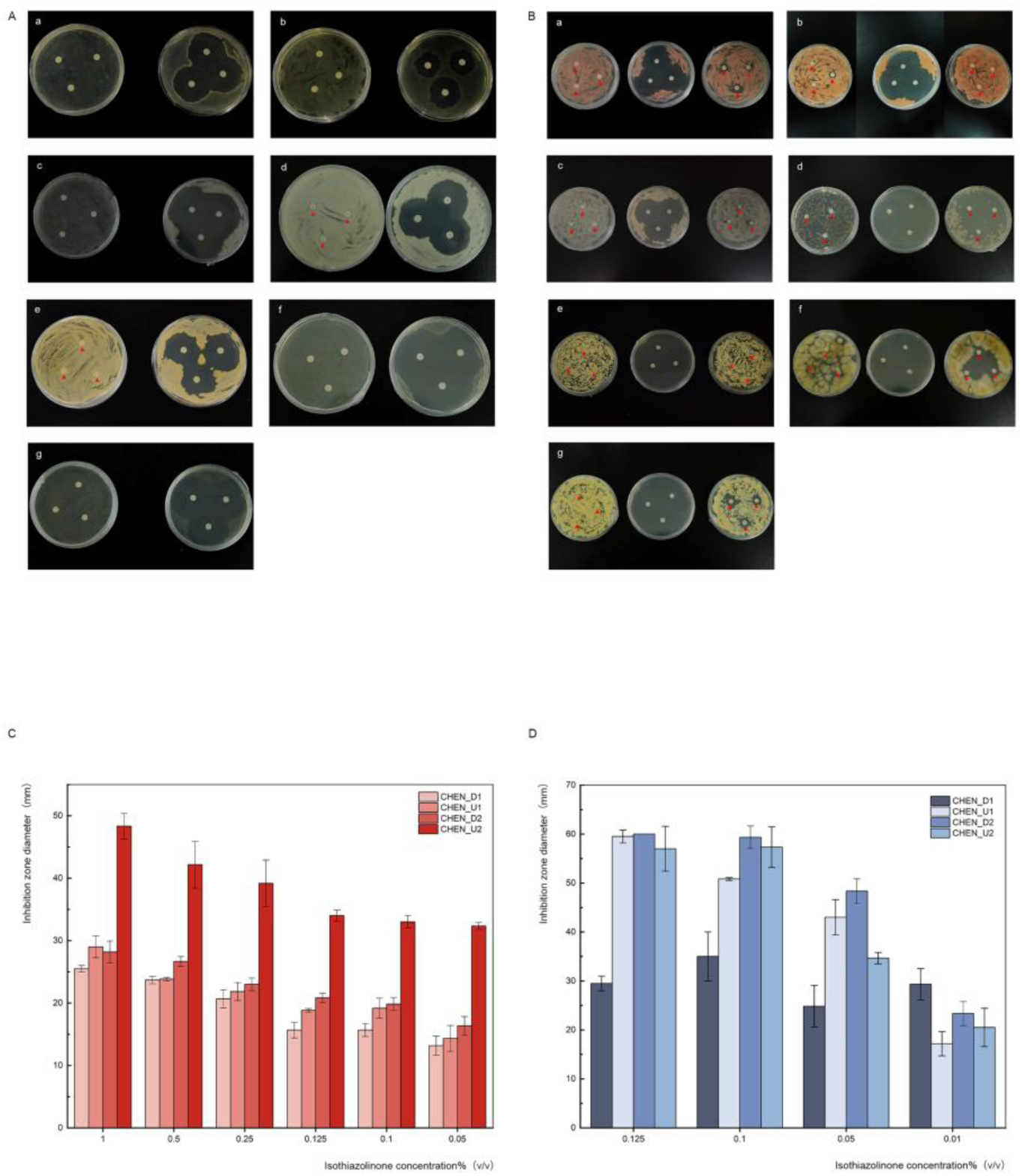
Antimicrobial sensitivity test of the microorganisms from Urb_CHEN. [A] Antimicrobial sensitivity test of the bacterial isolates using agar disk-diffusion method for two days. The left BPM plates were loaded with water; the right BPM plates were loaded with Isothiazolinone at a concentration of 1%(v/v). (a) *Pseudomonas* sp. (b) *Pantoea dispersa* strain. (c) *Paenibacillus* sp. (d) *Bacillus cereus* strain. (e) *Priestia megaterium* strain. (f) *Curtobacterium* sp. (g) *Microbacterium oleivorans* strain. [B] Antimicrobial sensitivity test of the fungal isolates using agar disk-diffusion method for three days. The PDA plates from left to right were loaded with water, Isothiazolinone at a concentration of 1% (v/v), and NaClO at a concentration of 1% (w/v). (a) *Rhodotorula mucilaginosa* strain. (b) *Cystobasidium* sp. (c) *Aureobasidium pullulans* strain. (d) *Cladosporium* sp. (e) *Trichoderma* sp. (f) *Daldinia* sp. (g) *Aspergillus sydowii* strain. The red triangles on both [a] and [b] indicate the location of partially indistinct disks. Three disks on the same plates were treated identically and used as a cross-reference. [C and D] Standard curve for the change of inhibition zone diameters using Isothiazolinone at a concentration gradient. Vertical lines indicate standard deviations. [c] Bacterial community for two days. [d] Fungal community for four days.

**Table 4:** The inhibition zones of bacteria (after two days) or fungi (after three days) were observed on BMP agar medium and PDA agar medium, respectively, using different biocides: Isothiazolinone at a concentration of 1% (v/v) and NaClO at a concentration of 1% (w/v)

| Isolates | Inhibition zone diameter (mm) |  |
| --- | --- | --- |
|  | Isothiazolinone | NaClO |
| CHEN_B1 | 33.73±0.76 | / |
| CHEN_B2 | 26.00±2.65 | / |
| CHEN_B3 | 45.33±1.15 | / |
| CHEN_B4 | 35.33±0.58 | / |
| CHEN_B5 | 32.00±1.00 | / |
| CHEN_B6 | 45.33±0.58 | / |
| CHEN_B7 | 42.00±2.00 | / |
| CHEN_F1 | 36.00±1.00 | 14.67±2.02 |
| CHEN_F2 | 23.83±1.04 | No inhibition |
| CHEN_F3 | 39.33±0.58 | No inhibition |
| CHEN_F4 | Complete inhibition | 44.67±2.31 |
| CHEN_F5 | Complete inhibition | 9.00±1.00 |
| CHEN_F6 | Complete inhibition | 27.33±1.15 |
| CHEN_F7 | Complete inhibition | 17.67±2.52 |

<sup>a</sup> Data are represented as mean ± SEM.
<sup>b</sup> The symbol '/' indicates that NaClO at a 1% (w/v) concentration did not show positive inhibitory effect on the bacterial isolates.

As Isothiazolinone exhibited the most potent antimicrobial effect, we focused our exploration on this biocide. Considering its potential environmental and human health risks at high concentrations, we aimed to reduce Isothiazolinone concentration while maintaining antimicrobial efficacy. We designed a concentration gradient experiment, preparing seven concentrations (1%, 0.5%, 0.25%, 0.125%, 0.1%, 0.05%, 0.01%). Although decreasing Isothiazolinone concentration led to a gradual reduction in bacterial antimicrobial sensitivity, even at 0.05%, noticeable inhibition zones persisted on bacteria community (Figure 5C), and most fungi community still exhibited significant inhibition at concentrations above 0.25%, except for CHEN_D1; and even at 0.01%, evident inhibition zones remained (Figure 5D). This extreme sensitivity of Chen Clan Ancestral Hall’s microbiota to Isothiazolinone is evident. Amplicon sequencing revealed a high *Epicoccum* abundance (35.61%) in CHEN_D1’s fungal community, possibly contributing to the limited antimicrobial effect of Isothiazolinone on CHEN_D1 and the irregular changes in antimicrobial sensitivity with decreasing concentrations.

## 4. Discussion

The amplicon sequencing results revealed a shared group of microorganisms on the damaged surfaces of ancestral halls. At the phylum level, the microbiota was dominated by *Firmicutes, Proteobacteria,* and *Ascomycota*; while at the genus level, it was primarily composed of *Bacillus, Pseudomonas, Paenibacillus, Acinetobacter, Toxicocladosporium, Cladosporium, Aspergillus,* and *Epicoccum*.

Among the microbiota derived from our samples, *Firmicutes, Proteobacteria, Bacteroidetes, Actinobacteria,* and *Ascomycota* are considered the main phyla for cellulose degradation (Liu et al., 2021). *Firmicutes* and *Proteobacteria*, key biofilm constituents, help retain vital nutrients and moisture for microorganisms(Ding et al., 2022). *Actinobacteria* colonization triggers pigment, organic acid, and polysaccharide production through secondary metabolism(Duan et al., 2017). This not only impacts the aesthetics of ancestral halls and causes wooden structure acid corrosion but also fosters biofilm formation on surfaces. In paint degradation, Phulpoto et al. discovered *Firmicutes* and *Actinobacteria* as dominant phyla, followed by *Proteobacteria*(Phulpoto et al., 2023). Comparing damaged and undamaged sites, damaged samples show increased abundance of *Firmicutes* and *Ascomycota*, while *Proteobacteria, Actinobacteria,* and *Phragmoplastophyta* prevail in undamaged areas. This distribution is hypothesized to stem from their distinct abilities in cellulose and paint degradation, as well as their varying environmental adaptability. And it should also be noted that changes in absolute abundances cannot be concluded from the relative abundance data(Li et al., 2020).

At the genus level, within our sample-derived microbiota, *Bacillus, Paenibacillus, Acinetobacter*, and *Pseudomonas* exhibit the ability to exclusively utilize lignin as a carbon source(Mendes et al., 2021; Xiong et al., 2020); *Aspergillus* and *Cladosporium*, isolated from lithographs, demonstrate cellulose-degrading capabilities (Coronado-Ruiz et al., 2018); *Pseudomonas*, *Cladosporium* and *Epicoccum* have been proven to be able to degrade paint as well as wood(Sanmartin et al., 2015; Shirakawa et al., 2002). In our study, *Cladosporium* shows higher abundance in damaged samples than undamaged ones. Isolated from Chen Clan Ancestral Hall, we confirmed its cellulose-degrading ability. It is also considered to be the main chromogen in artworks(Sabatini et al., 2018) and the main microbial disease of canoes preserved in the Oceanographic Museum of China(Zhang et al., 2019). Therefore, *Cladosporium* is likely to play a crucial role in ancestral hall deterioration. It is also suggested that using paint alone as the exclusive means to protect bare wood is not considered sufficient.

The KEGG analysis using PICRUSt reveals a potential comprehensive cellulose degradation process across all sampled surfaces. Microorganisms can acquire necessary nutrients and energy for survival via carbon fixation, nitrogen fixation, and the sulfur cycle. Similar to our study, Seward et al. employed PICRUSt to demonstrate heightened gene expression in purine/pyrimidine and amino-acid metabolism within mid-latitude peatlands(Seward et al., 2020). The method does have limitations as PICRUSt relies on predictions based on 16S rRNA genes rather than complete functional gene sequences, which could present a more precise prediction result. Wu et al. used metagenomics and metaproteomics analyses to identify microbial communities on the sandstone of Beishiku Temple in Northwest China and discovered key microbial groups involved in carbon, nitrogen, and sulfur cycles, as well as the potential specific pathways (Wu et al., 2023). However, the low biomass of our sampled wood surfaces hindered a more comprehensive exploration using a macrohistological approach. Future studies will be considered to identify the abundance of functional genes under optimized experimental conditions.

Simultaneously, we isolated eight bacterial and seven fungal strains from Chen Clan Ancestral Hall. Among the bacterial isolates, all seven aligned precisely with the primary bacterial group identified through amplicon sequencing. Three of the seven fungal isolates corresponded to the main fungal group. The alignment discrepancy in fungal isolates might be attributed to amplicon sequencing’s limitations in detecting less abundant species(Liu et al., 2018). In assessing the lignocellulose-degrading potential of these strains, six bacterial strains exhibited lignin-degrading activity, while five bacterial and five fungal strains demonstrated cellulose-degrading abilities. Notably, *Pantoea* has been observed to facilitate lignin degradation by producing laccase and lignin peroxidase (Atiwesh et al., 2022). *Microbacterium oleivorans* strain cannot utilize lignin and cellulose directly but may be able to utilize cellobiose for nutrients(Lian et al., 2016). While *Rhodotorula* and *Cystobacter* lack lignocellulose-degrading capabilities, their metabolic production of carotenoids could affect the aesthetic integrity of ancestral halls and shield biofilms against low-wavelength radiation (Cojoc et al., 2019).

Environmental factors can impact microbial diversity and distribution. For instance, the thriving cultural tourism industry introduces a microbiota through human activity, influencing microbial diversity on cultural relics. The microbial composition on ancestral hall surfaces, comprising prokaryotes like *Sedimentibacter*, *Trichococcus*, *Terribacillus*, *Pantoea*, *Proteiniphilum*, and eukaryotes such as *Rhizopus*, *Lichtheimia*, shows a positive correlation with visitor traffic. Studies indicate that *Trichococcus* dominates microbial communities in urban sewer systems(VandeWalle et al., 2012), while *Pantoea* can trace its source to human feces(Asai et al., 2019). *Rhizopus* and *Lichtheimia* are the most common fungi that cause mucormycosis(Gomes et al., 2011). Among the microorganisms isolated from Chen Clan Ancestral Hall, *Bacillus cereus* is often found on human hands (Adamczyk et al., 2020), and *Staphylococcus* is a bloodstream infection pathogen (Szczuka et al., 2015). Therefore, the microbial diversity on ancestral hall surfaces is likely influenced by contamination from frequent tourist activities.

New biocides, particularly plant essential oil extracts, have gained attention for their environmentally friendly nature(Cappitelli et al., 2020). However, in this study, cinnamon and thyme essential oils demonstrated weak antimicrobial effects. As for the mature commercial biocides we tested, Isothiazolinone proved more effective than sodium hypochlorite but poses environmental and human health risks when used excessively(Romani et al., 2022). Therefore, we aim to reduce the concentration of Isothiazolinone while ensuring a certain level of antimicrobial efficacy. Results show that Setting Isothiazolinone concentrations at 0.05% (v/v) for bacteria and 0.01% (v/v) for fungi in sensitivity tests on sample microbiota resulted in inhibition zone sizes exceeding 10 mm, with some surpassing 15 mm, which indicates moderate to high sensitivity(Fu et al., 2022). This suggests the feasibility of achieving antimicrobial effects while minimizing environmental impact through low-concentration Isothiazolinone use. However, caution regarding the harmful effects of Isothiazolinone remains essential.

## 5. Conclusion

In summary, we analyzed the microbial community on the surfaces of nine ancestral halls in the subtropical monsoon climate of Guangdong Province’s coastal areas. Employing both conventional culturing methods and amplicon sequencing, we identified a core group of microorganisms, including *Bacillus* sp., *Pseudomonas* sp., *Paenibacillus* sp., *Acinetobacter* sp., *Toxicocladosporium* sp., *Cladosporium* sp., *Aspergillus* sp., and *Epicoccum* sp. A proportion of the species isolated from the samples exhibited lignin or cellulose degradation capacity. Comparison between damaged and undamaged sites revealed similarities in microbial composition but differences in abundance. For instance, *Firmicutes* and *Ascomycota* were significantly more abundant in damaged surfaces while *Proteobacteria* and *Actinobacteria* were more prevalent in undamaged samples. Notably, Cladosporium, widely present in both sites, emerged as the primary cellulose-degrading genus. Given the similar microbial compositions between damaged and undamaged sites, it is suggested that using paint alone as the exclusive means to protect bare wood is insufficient. Microorganisms present on ancestral hall surfaces, including prokaryotes like *Sedimentibacter*, *Trichococcus, Terribacillus, Pantoea, Proteiniphilum* and eukaryotes such as *Rhizopus, Lichtheimia*, were found to be positively correlated with visitor flow rate, signifying the potential impact of visitor flow rates on the microbiota composition of cultural relics. Our observations underscore the microbial community’s abundance and diversity as a result of collaborative interactions among diverse environmental factors. Moreover, our investigation confirmed the efficacy of low-concentration Isothiazolinone as an inhibitor, but its practical use requires caution due to potential environmental and human health impacts.

## Supporting information

revised supplementary material

## Data accessibility

Raw sequencing reads about 16S amplicon sequencing and 18S amplicon sequencing are deposited at the NCBI Sequence Read Archive under the BioProject PRJNA925150 (SRR23110177 to SRR23110205) and PRJNA925313 (SRR23117924 to SRR23117952). DNA sequences of the bacterial and fungal isolates are deposited at the NCBI GenBank (accession numbers: OQ283719 to OQ283726, and OQ283737 to OQ283743). The datasets supporting this article have been uploaded as part of the supplemental material.

## Author’s contributions

**Ma.L.:** methodology, validation, formal analysis, investigation, resources, data curation, visualization, and writing – original draft; **X.S.:** methodology, validation, formal analysis, investigation, resources, data curation, visualization, and writing – original draft; **Q.W.:** investigation, visualization, and writing – original draft; **T.Y.:** investigation and resources; **Mu.L.:** investigation, and resources; **P.A.:** investigation; **J.R.L.:** conceptualization, validation, formal analysis, writing – review and editing, supervision, and funding acquisition; **J.L.:** conceptualization, validation, formal analysis, writing – review and editing, and funding acquisition.

## Ethics

Permission was obtained from the Villager Committees and staff in charge of the ancestral halls before sampling was carried out.

## Acknowledgments

We thank Zhen Yi, Haigan Huang, and all the ancestral hall conservators for their help in collecting microbiological samples, Yueyue Zhu and Zhenyang Zhang for lab work assistance, the Guangdong Folk Art Museum, the Kongmei Villager Committee and the Lingnan Impression Garden for support in the project, ChatGPT for language quality improvements.

