## Supplementary material for "Insights into the Microbial Weathering of Chinese Wooden Ancestral Halls in Guangdong Province": revised supplementary material

### Supplementary Figures and Tables

#### Supplementary Figures

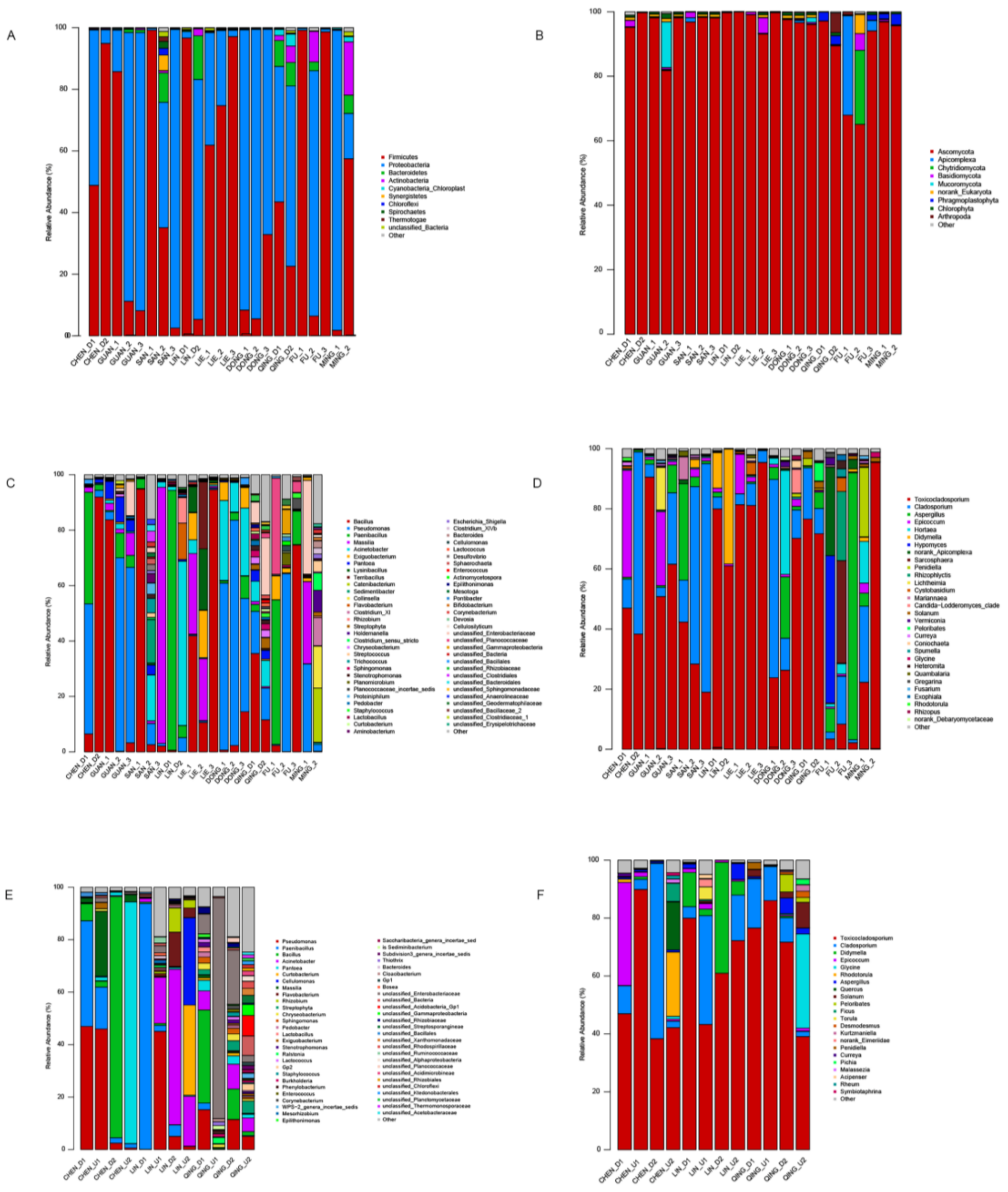


**Supplementary Figure 1. Taxonomic analysis of microbial communities in the samples.**

Percentage of amplicon reads mapped to each sample of all ancestral halls.

(A, B, C and D) Phylum-level taxonomy and genus-level taxonomy of the microbiome in the 26 samples collected from damaged surfaces of nine ancestral halls with varying visitor flow rates under the same climate condition. The taxonomic analysis was performed using 16S amplicon sequencing for (A)bacterial phyla and (C)bacterial genera, and 18S amplicon sequencing for (B)fungal phyla and (D)fungal genera.

(E and F) Genus-level taxonomy of the microbiome in the samples collected from damaged and undamaged surfaces of Chen Clan Ancestral Hall, Lin Clan Ancestral Hall, and Qingxi Ancestral Hall. The taxonomic analysis was conducted using 16S amplicon sequencing for (E)bacterial genera and 18S amplicon sequencing for (F)fungal genera.

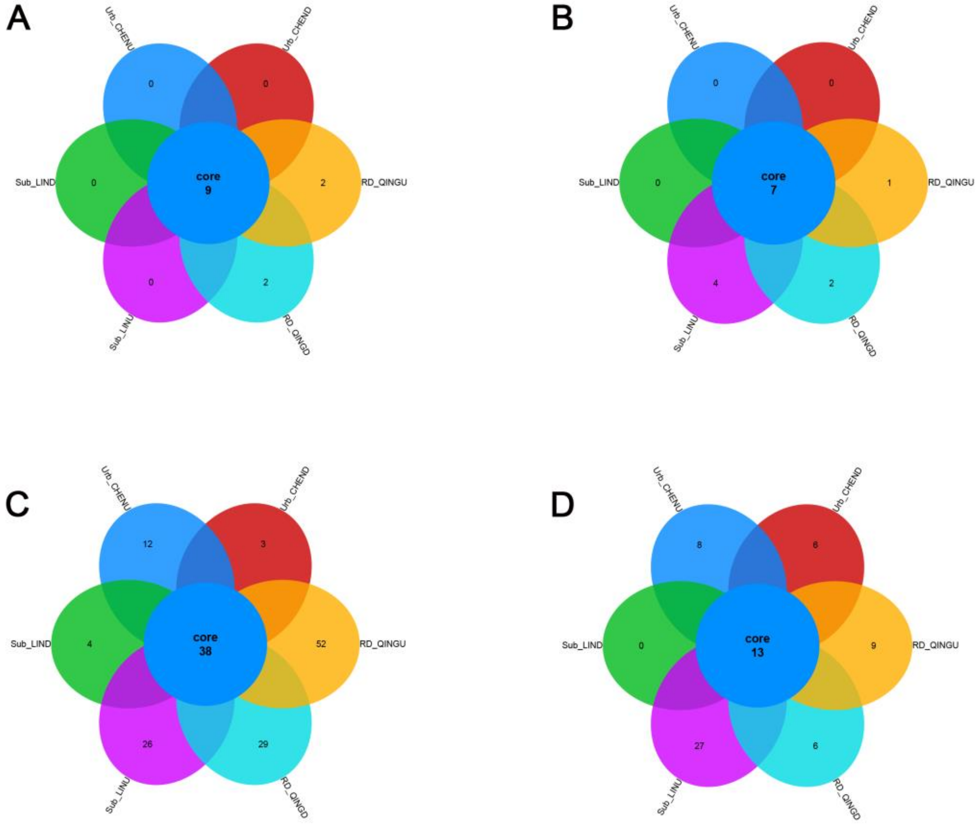


**Supplementary Figure 2. The unique and shared (A) Bacterial phyla (B) Fungal phyla (C) Bacterial genera (D) Fungal genera in damaged and undamaged surface from three ancestral halls** (Urb_CHEND and Urb_CHENU in Chen Clan Ancestral Hall, Sub_LIND and Sub_LINU in Lin Clan Ancestral Hall, RD_QINGD and RD_QINGU in Qingxi Ancestral Hall).

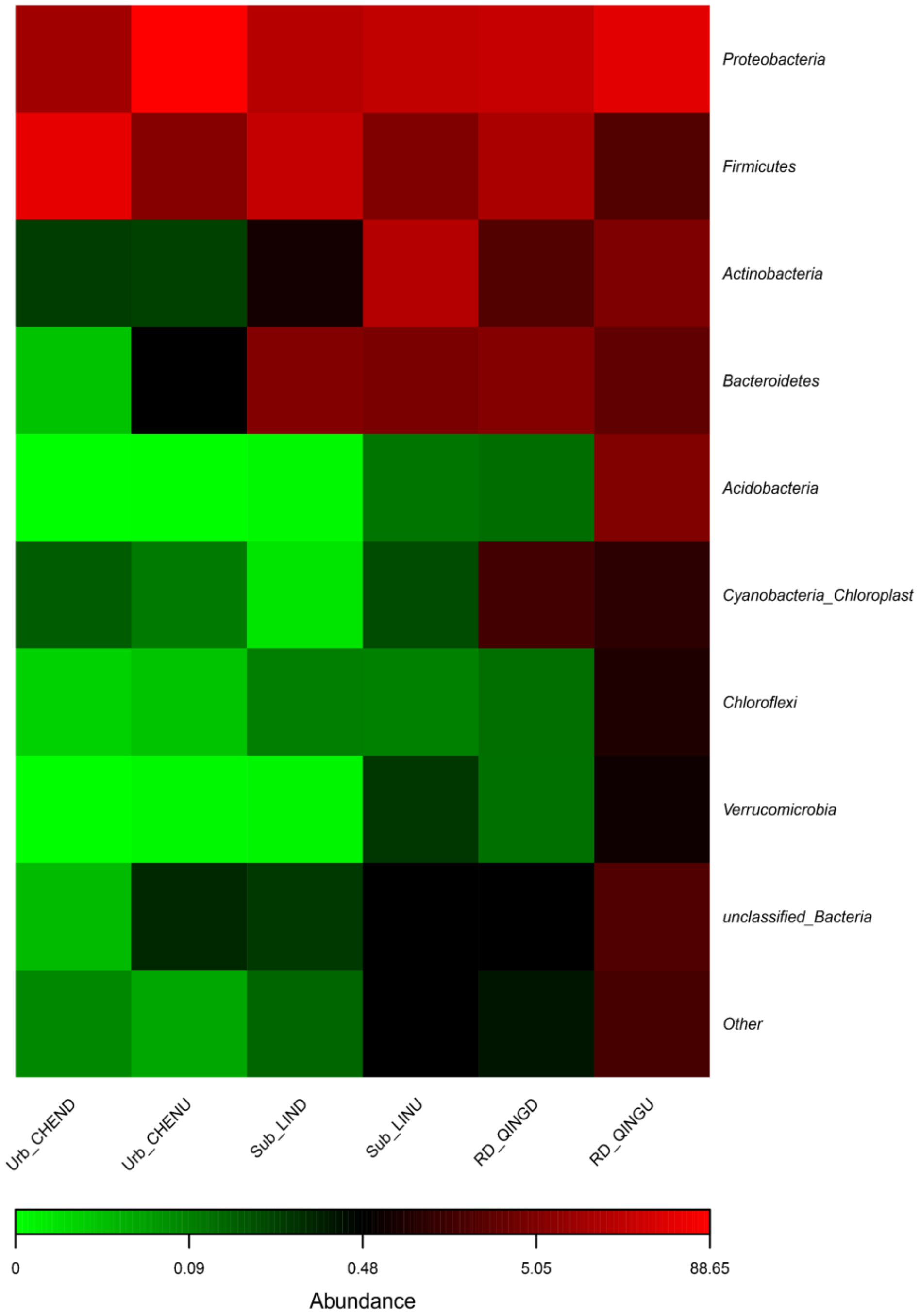


**Supplementary Figure 3. Heatmap showing the diversity of bacteria at the phylum level in different groups.**

Color intensity signifies abundance ranging from green (low abundance), to black (medium abundance) and to red (high abundance).

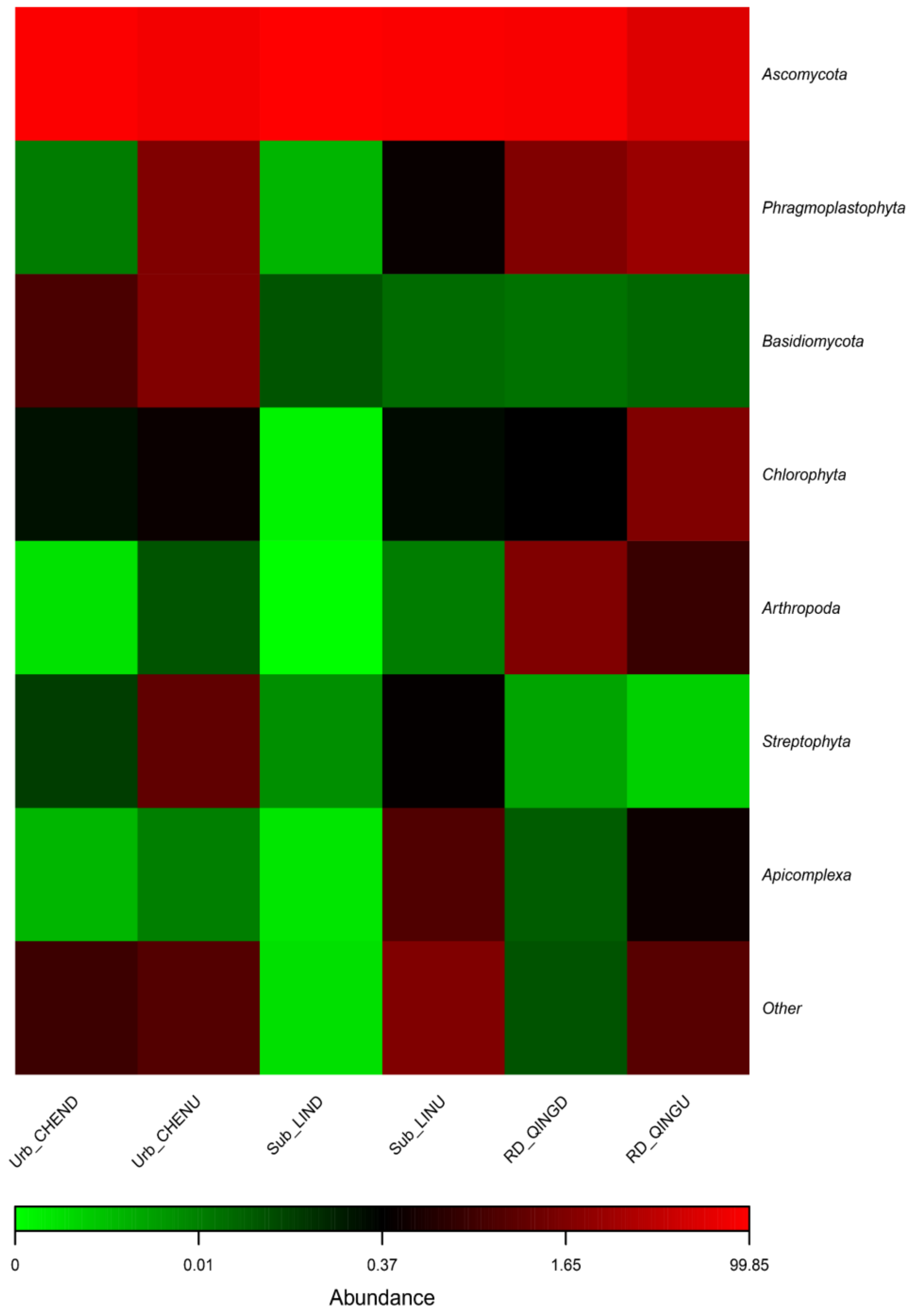


**Supplementary Figure 4. Heatmap showing the diversity of fungi at the phylum level in different groups.**

Color intensity signifies abundance ranging from green (low abundance), to black (medium abundance) and to red (high abundance).

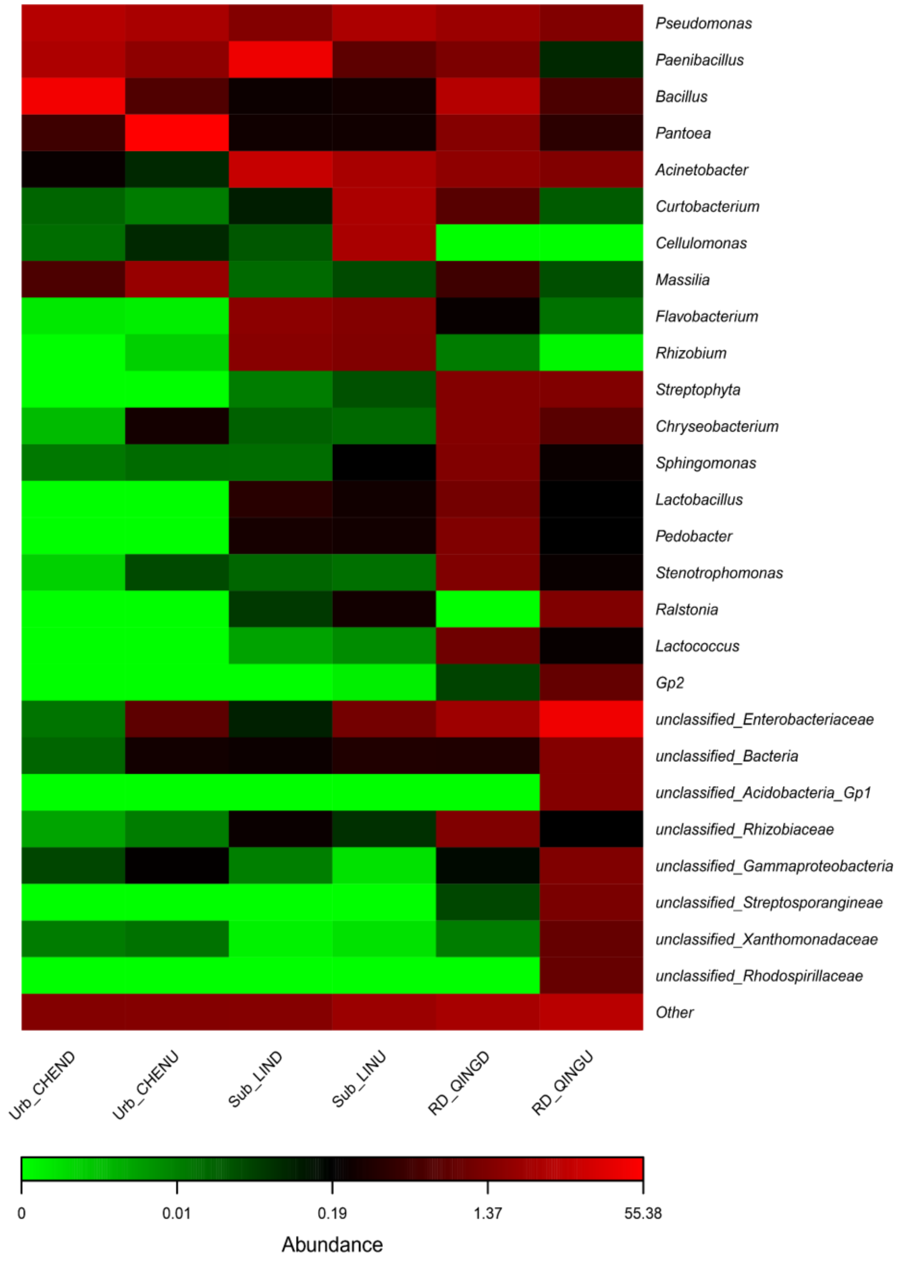


**Supplementary Figure 5. Heatmap showing the diversity of bacteria at the genus level in different groups.**

Color intensity signifies abundance ranging from green (low abundance), to black (medium abundance) and to red (high abundance).

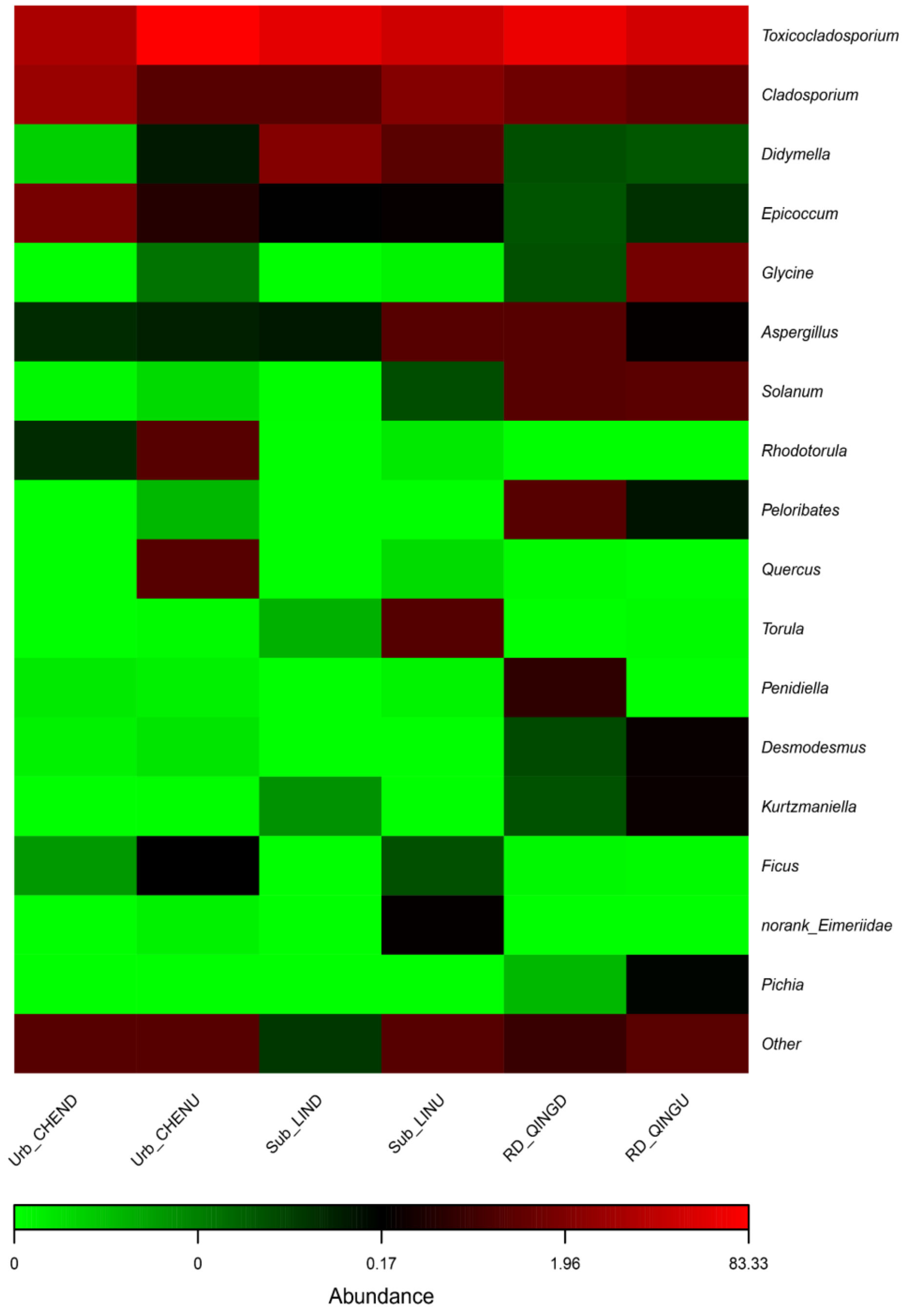


**Supplementary Figure 6. Heatmap shows the diversity of fungi at the genus level in different groups.**

Color intensity signifies abundance ranging from green (low abundance), to black (medium abundance) and to red (high abundance).

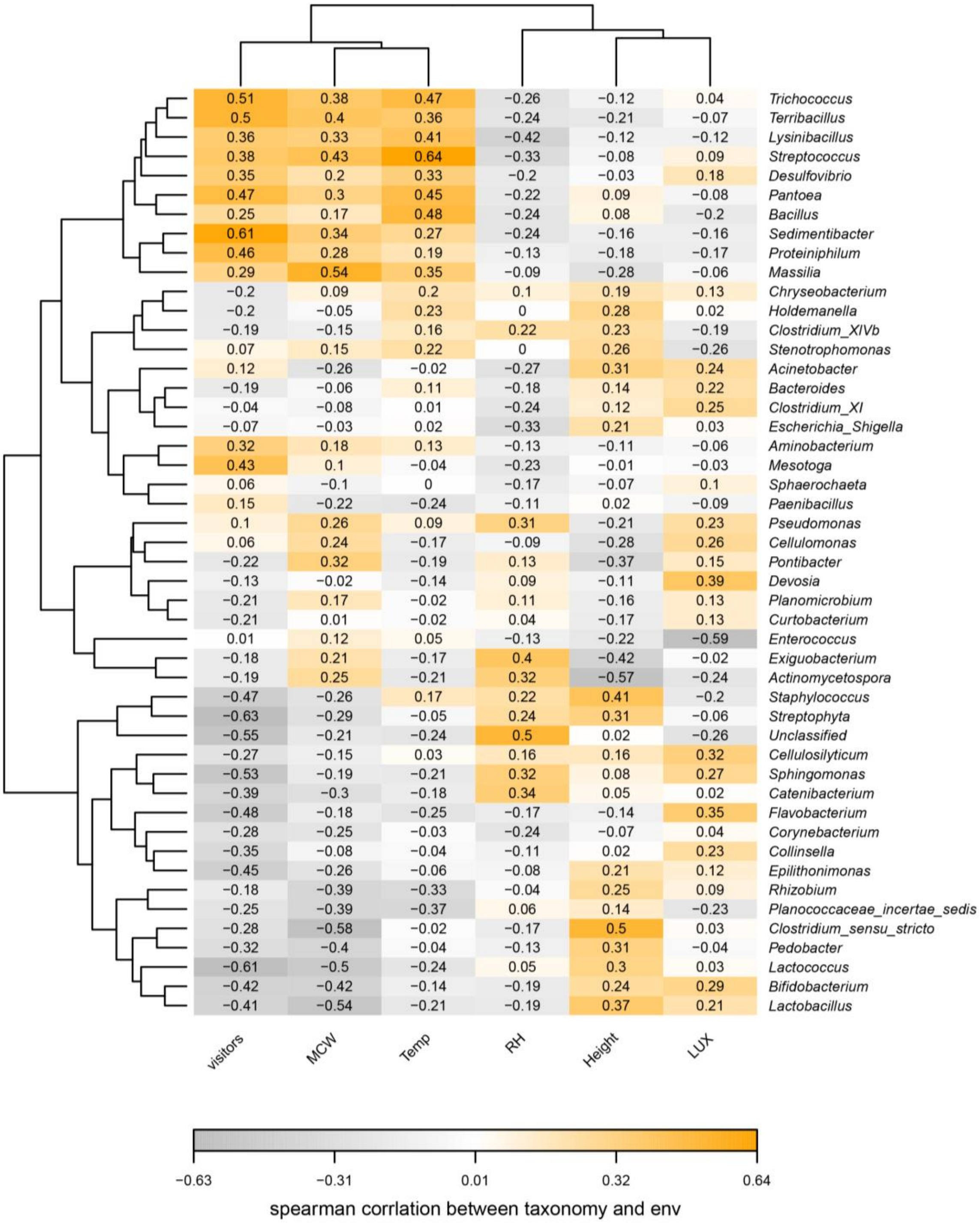


**Supplementary Figure 7. Correlation heatmap of environmental factors with the microbial community on prokaryotic genus level.**

The color key for the correlation values is displayed in the bottom panel inset. Positive correlations are indicated by orange text, negative correlations are represented in grey, and non-significant correlations are shown in white. The values represent the Spearman's rank coefficient of correlation between each prokaryotic genus and each environmental factor.

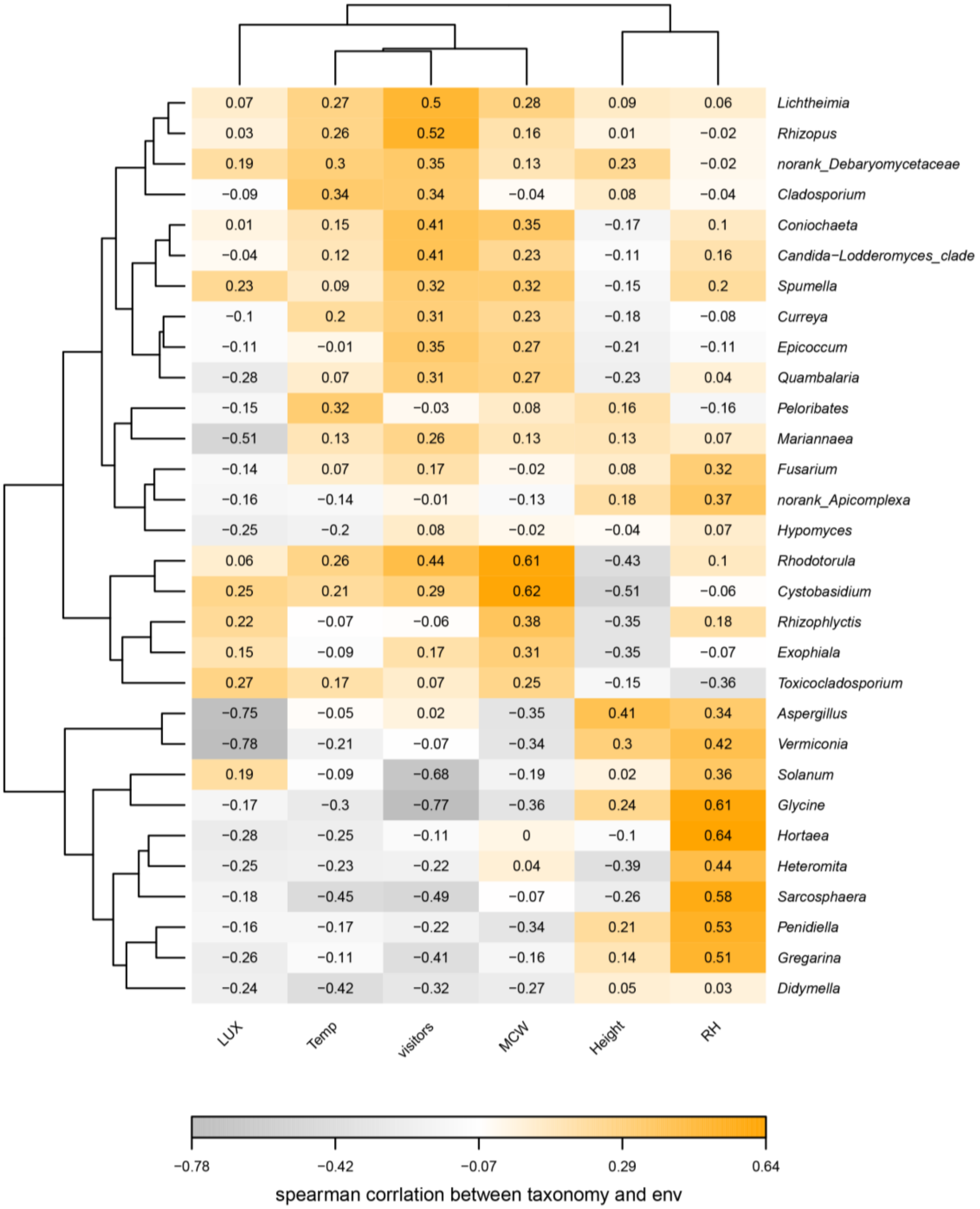


**Supplementary Figure 8. Correlation heatmap of environmental factors with the microbial community on eukaryotic genus level.**

The color key for the correlation values is displayed in the bottom panel inset. Positive correlations are indicated by orange text, negative correlations are represented in grey, and non-significant correlations are shown in white. The values represent the Spearman's rank coefficient of correlation between each prokaryotic genus and each environmental factor.

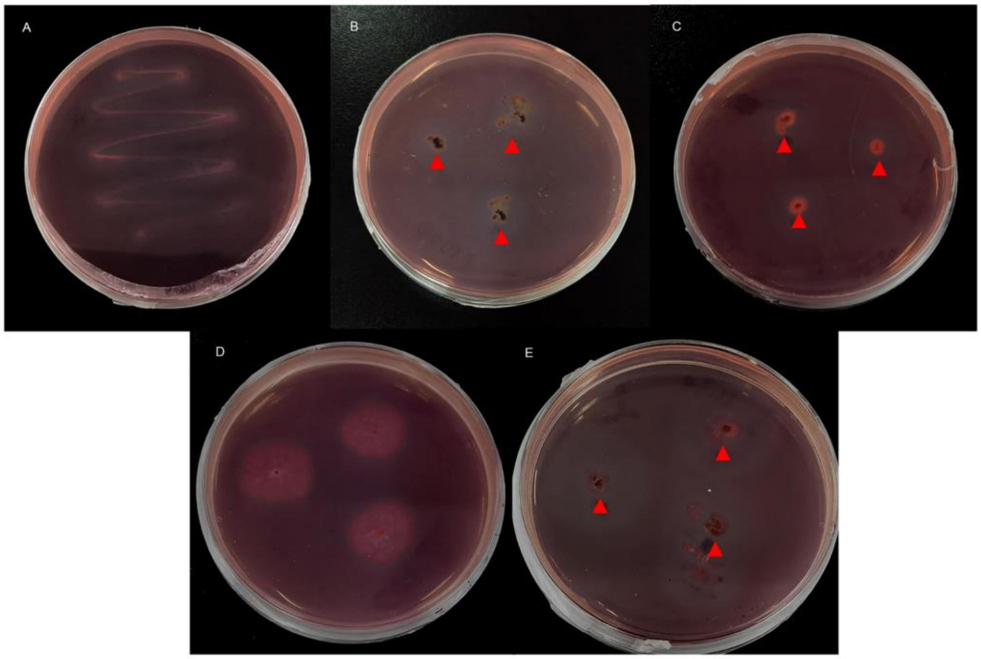


**Supplementary Figure 9. Verification of the degrading ability of microorganisms from Urb_CHEN based on conventional culturing methods**

Hydrolytic zones of the fungal isolates on CMC-degrading agar medium after three days of incubation. (A) *Aureobasidium pullulans* strain. (B) *Cladosporium* sp. (C) *Trichoderma* sp. (D) *Daldinia* sp. (E) *Aspergillus sydowii* strain.

The plate streaking method instead of inoculation was employed to assess cellulose-degradation ability due to the viscous nature of the colony of (A)*Aureobasidium pullulans* strain.

### Supplementary Tables

#### Table S1. The associated environmental factor information from samples

| Sample | Temperature (℃) | Relative humidity (%) | Light (Lux) | The moisture contents of the wood (%) | Height (cm) | Visitor flow rate |
|---|---|---|---|---|---|---|
| CHEN_D1 and CHEN_U1 | 35.4 | 56.7 | 2168 | 18.5 | 10 | +++ |
| CHEN_D2 and CHEN_U2 | 35.0 | 59.5 | 2085 | 12.5 | 50 | +++ |
| GUAN_1 | 34.2 | 70.9 | 512 | 21 | 73 | +++ |
| GUAN_2 | 33.7 | 70.9 | 2478 | 28 | 80 | +++ |
| GUAN_3 | 33.4 | 73.5 | 33.4 | 14.5 | 23 | +++ |
| SAN_1 | 33.9 | 65.9 | 602 | 11 | 283 | +++ |
| SAN_2 | 34.5 | 62.8 | 482 | 8.5 | 287 | +++ |
| SAN_3 | 34.4 | 63.5 | 1393 | 12 | 127 | +++ |
| LIN_D1 and LIN_U1 | 27.5 | 56.4 | 727 | 90 | 4.5 | ++ |
| LIN_D2 and LIN_U2 | 27.8 | 54.9 | 4137 | 43 | 9.5 | ++ |
| LIE_1 | 31.9 | 66.8 | 633 | 30 | 14.5 | ++ |
| LIE_2 | 31.9 | 67.2 | 1402 | 40 | 14.5 | ++ |
| LIE_3 | 32.0 | 67.5 | 1287 | 12 | 14.5 | ++ |
| DONG_1 | 31.0 | 79.3 | 1453 | 65 | 4 | ++ |
| DONG_2 | 31.3 | 78.5 | 559 | 78 | 10.5 | ++ |
| DONG_3 | 31.6 | 75.3 | 2422 | 102 | 6.5 | ++ |
| QING_D1 and QING_U1 | 33.5 | 70.8 | 7512 | 128 | 5.5 | + |
| QING_D2 and QING_U2 | 34.1 | 69.6 | 466 | 69 | 12.5 | + |
| FU_1 | 30.1 | 82.4 | 128 | 51 | 5.5 | + |
| FU_2 | 31.0 | 82.0 | 2765 | 13 | 16 | + |
| FU_3 | 31.5 | 81.0 | 329 | 153 | 6 | + |
| MING_1 | 30.8 | 83.0 | 252 | 67.5 | 8 | + |
| MING_2 | 32.1 | 78.3 | 6658 | 93 | 8 | + |
| MING_3 | 31.4 | 80.7 | 257 | 4.0 | 67.5 | + |

[a] The symbol "+" denotes hourly visitor calculations of 10, "++" denotes hourly visitor calculations of 100, and "+++" denotes hourly visitor calculations of 10,000.

**Table S2. Alpha diversity of all samples from 16S amplicon sequencing**

| Sample | Number | OTUs | Shannon | Chao | Ace | Simpson | Shannon evenness |
|---|---|---|---|---|---|---|---|
| CHEN_D1 | 27156 | 157 | 1.35 | 245.50 | 235.54 | 0.38 | 0.27 |
| CHEN_U1 | 32587 | 171 | 2.02 | 202.94 | 212.72 | 0.26 | 0.39 |
| CHEN_D2 | 28779 | 116 | 0.57 | 198.50 | 284.95 | 0.82 | 0.12 |
| CHEN_U2 | 48786 | 174 | 0.54 | 235.03 | 245.50 | 0.84 | 0.11 |
| GUAN_1 | 30398 | 152 | 1.52 | 228.96 | 287.96 | 0.42 | 0.30 |
| GUAN_2 | 20876 | 177 | 1.99 | 262.95 | 293.49 | 0.33 | 0.38 |
| GUAN_3 | 25352 | 170 | 1.92 | 235.36 | 242.41 | 0.23 | 0.37 |
| SAN_1 | 86467 | 133 | 0.33 | 247.00 | 270.10 | 0.89 | 0.07 |
| SAN_2 | 24060 | 267 | 3.80 | 297.75 | 302.84 | 0.06 | 0.68 |
| SAN_3 | 60696 | 165 | 0.49 | 200.65 | 199.79 | 0.86 | 0.10 |
| LIN_D1 | 50237 | 116 | 0.75 | 124.27 | 124.93 | 0.78 | 0.16 |
| LIN_U1 | 52349 | 78 | 0.24 | 81.60 | 82.13 | 0.94 | 0.05 |
| LIN_D2 | 49986 | 120 | 2.14 | 120.40 | 121.60 | 0.20 | 0.45 |
| LIN_U2 | 50453 | 118 | 1.84 | 120.33 | 121.40 | 0.25 | 0.39 |
| DONG_1 | 32508 | 181 | 1.57 | 229.36 | 234.05 | 0.28 | 0.30 |
| DONG_2 | 41988 | 193 | 1.18 | 240.90 | 242.53 | 0.53 | 0.22 |
| DONG_3 | 36795 | 176 | 2.01 | 224.36 | 233.69 | 0.18 | 0.39 |
| LIE_1 | 29620 | 168 | 2.27 | 207.52 | 219.56 | 0.15 | 0.44 |
| LIE_2 | 28759 | 185 | 2.13 | 250.21 | 252.13 | 0.16 | 0.41 |
| LIE_3 | 29620 | 144 | 0.49 | 189.37 | 204.18 | 0.87 | 0.10 |
| QING_D1 | 59240 | 151 | 2.91 | 152.00 | 152.03 | 0.17 | 0.58 |
| QING_U1 | 71339 | 97 | 0.15 | 114.00 | 105.82 | 0.97 | 0.03 |
| QING_D2 | 51303 | 226 | 3.86 | 227.50 | 228.94 | 0.06 | 0.71 |
| QING_U2 | 62297 | 472 | 5.41 | 472.00 | 472.18 | 0.01 | 0.88 |
| FU_1 | 72023 | 104 | 1.26 | 153.40 | 182.69 | 0.38 | 0.27 |
| FU_2 | 55620 | 140 | 1.86 | 148.27 | 150.46 | 0.42 | 0.38 |
| FU_3 | 78882 | 95 | 1.29 | 99.00 | 98.38 | 0.51 | 0.28 |
| MING_1 | 47945 | 99 | 1.80 | 116.77 | 116.29 | 0.22 | 0.39 |
| MING_2 | 52721 | 389 | 3.61 | 394.00 | 389.67 | 0.08 | 0.60 |

**Table S3. Alpha diversity of all samples from 18S amplicon sequencing**

| Sample | Number | OTUs | Shannon | Chao | Ace | Simpson | Shannon evenness |
|---|---|---|---|---|---|---|---|
| CHEN_D1 | 43781 | 102 | 1.60 | 119.10 | 115.07 | 0.27 | 0.35 |
| CHEN_U1 | 47405 | 97 | 0.79 | 110.91 | 109.13 | 0.68 | 0.17 |
| CHEN_D2 | 44674 | 18 | 0.47 | 18.60 | 19.57 | 0.75 | 0.16 |
| CHEN_U2 | 21523 | 24 | 0.29 | 27.00 | 30.14 | 0.90 | 0.09 |
| GUAN_1 | 47013 | 88 | 0.64 | 97.55 | 97.90 | 0.75 | 0.14 |
| GUAN_2 | 32805 | 108 | 1.89 | 118.56 | 121.35 | 0.23 | 0.40 |
| GUAN_3 | 39443 | 94 | 0.88 | 102.57 | 103.96 | 0.63 | 0.19 |
| SAN_1 | 45615 | 17 | 1.15 | 18.00 | 22.68 | 0.39 | 0.41 |
| SAN_2 | 55131 | 100 | 0.79 | 142.17 | 114.17 | 0.70 | 0.17 |
| SAN_3 | 50142 | 98 | 0.43 | 119.43 | 117.11 | 0.86 | 0.09 |
| LIN_D1 | 107730 | 18 | 0.65 | 18.00 | 18.45 | 0.70 | 0.23 |
| LIN_U1 | 84446 | 71 | 1.08 | 71.00 | 71.00 | 0.64 | 0.25 |
| LIN_D2 | 98860 | 11 | 0.70 | 12.50 | 13.43 | 0.52 | 0.29 |
| LIN_U2 | 118000 | 25 | 0.59 | 28.00 | 31.53 | 0.74 | 0.18 |
| LIE_1 | 45769 | 33 | 0.79 | 36.33 | 35.63 | 0.63 | 0.23 |
| LIE_2 | 35272 | 85 | 1.10 | 89.40 | 91.69 | 0.54 | 0.25 |
| LIE_3 | 53975 | 20 | 0.63 | 20.33 | 21.14 | 0.60 | 0.21 |
| DONG_1 | 35043 | 94 | 0.66 | 106.67 | 107.42 | 0.75 | 0.15 |
| DONG_2 | 42302 | 101 | 1.31 | 124.75 | 115.70 | 0.39 | 0.28 |
| DONG_3 | 50920 | 100 | 1.55 | 109.55 | 108.32 | 0.33 | 0.34 |
| QING_D1 | 110079 | 40 | 0.41 | 40.00 | 40.33 | 0.86 | 0.11 |
| QING_U1 | 77244 | 34 | 0.18 | 36.00 | 36.93 | 0.95 | 0.05 |
| QING_D2 | 58217 | 46 | 1.07 | 46.00 | 47.11 | 0.62 | 0.28 |
| QING_U2 | 86593 | 46 | 1.82 | 46.00 | 47.00 | 0.28 | 0.48 |
| FU_1 | 101982 | 49 | 1.52 | 51.50 | 51.47 | 0.33 | 0.39 |
| FU_2 | 83946 | 62 | 1.95 | 64.33 | 67.79 | 0.21 | 0.47 |
| FU_3 | 75404 | 40 | 0.65 | 41.00 | 41.43 | 0.78 | 0.18 |
| MING_1 | 62146 | 67 | 1.72 | 67.00 | 67.19 | 0.29 | 0.41 |
| MING_2 | 77446 | 21 | 0.27 | 21.00 | 21.00 | 0.91 | 0.09 |

**Table S4. The dominant prokaryotic phyla on the damaged surfaces of nine ancestral halls (%)**

| | Urb_CHEN | Urb_GUAN | Urb_SAN | Sub_LIN | Sub_LIE | Sub_DONG | RD_QING | RD_FU | RD_MING |
|---|---|---|---|---|---|---|---|---|---|
| *Firmicutes* | 72.36 | 38.20 | 55.87 | 50.19 | 77.99 | 15.45 | 33.27 | 74.60 | 31.20 |
| *Proteobacteria* | 26.94 | 60.51 | 40.40 | 40.65 | 20.88 | 83.98 | 51.06 | 21.47 | 53.71 |
| *Bacteroidetes* | 0.04 | 0.66 | 1.38 | 7.36 | 0.21 | 0.14 | 8.02 | 0.81 | 3.20 |
| *Actinobacteria* | 0.29 | 0.24 | 0.15 | 1.21 | 0.13 | 0.13 | 3.46 | 2.67 | 9.25 |
| *Cyanobacteria_Chloroplast* | 0.20 | 0.17 | 0.11 | 0.02 | 0.19 | 0.09 | 2.87 | 0.12 | 1.08 |
| Other | 0.16 | 0.22 | 2.09 | 0.58 | 0.60 | 0.21 | 1.32 | 0.33 | 1.57 |

**Table S5. The dominant eukaryotic phyla on the damaged surfaces of nine ancestral halls (%)**

| | Urb_CHEN | Urb_GUAN | Urb_SAN | Sub_LIN | Sub_LIE | Sub_DONG | RD_QING | RD_FU | RD_MING |
|---|---|---|---|---|---|---|---|---|---|
| *Ascomycota* | 97.37 | 93.68 | 97.75 | 99.85 | 97.79 | 96.66 | 94.48 | 74.53 | 96.24 |
| *Apicomplexa* | 0.01 | 0.05 | 0.42 | 0 | 0 | 0.23 | 0.11 | 12.98 | 0.08 |
| *Chytridiomycota* | 0.11 | 0.21 | 0.13 | 0 | 0.06 | 0.24 | 0 | 7.40 | 0 |
| *Phragmoplastophyta* | 0.02 | 0.01 | 0.02 | 0.01 | 0.01 | 0.01 | 2.74 | 0.63 | 2.58 |
| *Basidiomycota* | 1.11 | 0.22 | 0.58 | 0.13 | 1.62 | 0.30 | 0.05 | 1.71 | 0.23 |
| *norank_Eukaryota* | 0.47 | 0.82 | 0.54 | 0 | 0.25 | 0.92 | 0.02 | 1.93 | 0.22 |
| *Mucoromycota* | 0.04 | 3.93 | 0.05 | 0 | 0.02 | 0.68 | 0 | 0 | 0.02 |
| *Arthropoda* | 0 | 0 | 0 | 0 | 0 | 0.03 | 2.10 | 0.38 | 0.06 |
| Other | 0.89 | 1.07 | 0.51 | 0.01 | 0.26 | 0.94 | 0.50 | 0.45 | 0.57 |

**Table S6. The dominant prokaryotic genera on the damaged surfaces of nine ancestral halls (%)**

| | Urb_CHEN | Urb_GUAN | Urb_SAN | Sub_LIN | Sub_LIE | Sub_DONG | RD_QING | RD_FU | RD_MING |
|---|---|---|---|---|---|---|---|---|---|
| *Bacillus* | 50.13 | 32.26 | 48.95 | 0.30 | 49.41 | 5.87 | 23.80 | 29.45 | 0.17 |
| *Pseudomonas* | 24.22 | 43.06 | 1.51 | 2.63 | 0.43 | 61.73 | 13.36 | 17.08 | 16.18 |
| *Paenibacillus* | 20.73 | 4.50 | 1.80 | 48.22 | 0.24 | 3.93 | 1.34 | 23.10 | 0.06 |
| *Massilia* | 0.89 | 3.68 | 32.93 | 0.04 | 17.22 | 0.21 | 0.77 | 0.07 | 13.96 |
| *Acinetobacter* | 0.27 | 1.23 | 3.11 | 30.87 | 1.96 | 20.51 | 8.35 | 0.13 | 0.35 |
| *Exiguobacterium* | 0.14 | 0.42 | 0.04 | 0.01 | 9.03 | 4.40 | 0.23 | 3.51 | 0.72 |
| *Pantoea* | 0.76 | 6.70 | 0.67 | 0.34 | 0.23 | 0.31 | 3.61 | 0.02 | 0.12 |
| *Lysinibacillus* | 0.09 | 0.07 | 0.12 | 0.03 | 10.55 | 0.03 | 0.07 | 0.01 | 0.02 |
| *Catenibacterium* | 0 | 0 | 0 | 0 | 0 | 0 | 0 | 0 | 10.30 |
| *Terribacillus* | 0.16 | 0.10 | 0.03 | 0 | 8.06 | 0.04 | 0 | 0.01 | 0 |
| *Collinsella* | 0 | 0 | 0 | 0 | 0 | 0 | 0.07 | 0 | 8.00 |
| *Flavobacterium* | 0 | 0 | 0 | 6.79 | 0 | 0 | 0.25 | 0.07 | 0.08 |
| *Clostridium_XI* | 0 | 0 | 0 | 0 | 0 | 0 | 0 | 0 | 5.33 |
| *Rhizobium* | 0 | 0 | 0 | 4.77 | 0 | 0 | 0.02 | 0 | 0.30 |
| *Holdemanella* | 0 | 0 | 0 | 0 | 0 | 0 | 0.03 | 0 | 4.15 |
| *Streptophyta* | 0 | 0 | 0.02 | 0.01 | 0 | 0 | 2.86 | 0.10 | 0.98 |
| *Clostridium_sensu_stricto* | 0.01 | 0 | 0.11 | 0.05 | 0.01 | 0.01 | 0.16 | 0.02 | 3.31 |
| *Chryseobacterium* | 0.01 | 0.37 | 0.01 | 0.05 | 0 | 0.01 | 2.55 | 0.02 | 0.21 |
| *Streptococcus* | 0.09 | 0.07 | 0.25 | 0.01 | 0.09 | 0.04 | 0.06 | 0 | 2.15 |
| *Sphingomonas* | 0.02 | 0.03 | 0.02 | 0.04 | 0.02 | 0.03 | 2.23 | 0.21 | 0.08 |
| *Sedimentibacter* | 0.02 | 0.06 | 2.23 | 0.01 | 0.17 | 0.07 | 0 | 0 | 0 |
| *Stenotrophomonas* | 0 | 0.36 | 0.22 | 0.05 | 0.01 | 0.01 | 1.48 | 0.02 | 0.01 |
| *Lactobacillus* | 0 | 0 | 0.05 | 0.56 | 0 | 0 | 1.26 | 0.01 | 0.19 |
| *Pedobacter* | 0 | 0.19 | 0 | 0.39 | 0 | 0 | 1.40 | 0.01 | 0.02 |
| *Planococcaceae_incertae_sedis* | 0 | 0 | 0 | 0.02 | 0 | 0 | 0.01 | 1.64 | 0 |
| *Bacteroides* | 0 | 0 | 0 | 0 | 0 | 0 | 0.03 | 0 | 1.35 |
| *Clostridium_XlVb* | 0 | 0 | 0.01 | 0 | 0 | 0 | 0.20 | 0 | 1.17 |
| *Planomicrobium* | 0 | 0.06 | 0 | 0 | 0 | 0 | 0.12 | 1.18 | 0 |
| *Lactococcus* | 0 | 0 | 0 | 0.01 | 0 | 0 | 1.22 | 0 | 0.03 |
| *unclassified_Enterobacteriaceae* | 0.02 | 3.78 | 0.31 | 0.14 | 0.04 | 0.11 | 13.93 | 0.03 | 16.05 |
| *unclassified_Planococcaceae* | 0 | 0 | 0 | 0.01 | 0 | 0 | 0.92 | 13.84 | 0.07 |
| *unclassified_Gammaproteobacteria* | 0.09 | 0.62 | 0.06 | 0.01 | 0.08 | 0.34 | 0.18 | 2.29 | 0.60 |
| *unclassified_Rhizobiaceae* | 0.01 | 0.02 | 0.01 | 0.29 | 0.01 | 0.01 | 1.90 | 0.07 | 0.25 |
| Other | 2.32 | 2.42 | 7.52 | 4.34 | 2.42 | 2.32 | 17.56 | 7.11 | 13.80 |

### Table S7. The dominant eukaryotic genera on the damaged surfaces of nine ancestral halls (%)

| | Urb_CHEN | Urb_GUAN | Urb_SAN | Sub_LIN | Sub_LIE | Sub_DONG | RD_QING | RD_FU | RD_MING |
|---|---|---|---|---|---|---|---|---|---|
| *Toxicocladosporium* | 42.79 | 70.45 | 29.80 | 70.84 | 86.77 | 44.02 | 74.90 | 4.66 | 62.63 |
| *Cladosporium* | 34.35 | 10.16 | 50.17 | 2.08 | 4.72 | 24.73 | 14.06 | 6.32 | 11.36 |
| *Aspergillus* | 0.60 | 3.74 | 11.63 | 0.82 | 0.18 | 8.05 | 2.26 | 28.65 | 1.94 |
| *Epicoccum* | 18.39 | 7.44 | 1.33 | 1.04 | 5.02 | 1.55 | 0.17 | 0.45 | 1.53 |
| *Didymella* | 0.05 | 0.11 | 1.01 | 24.55 | 0.09 | 0.07 | 0.23 | 0.46 | 0.68 |
| *Hortaea* | 0.18 | 0.43 | 0.27 | 0 | 0.10 | 11.75 | 0.14 | 1.57 | 6.21 |
| *Hypomyces* | 0.01 | 0.02 | 0.01 | 0.03 | 0.06 | 0 | 0 | 19.12 | 0.02 |
| *norank_Apicomplexa* | 0 | 0 | 0.42 | 0 | 0 | 0.15 | 0 | 12.36 | 0 |
| *Penidiella* | 0.02 | 0.02 | 0.02 | 0 | 0.01 | 0.52 | 1.55 | 0.05 | 10.33 |
| *Sarcosphaera* | 0 | 0 | 0 | 0 | 0 | 0 | 0 | 10.98 | 0.08 |
| *Rhizophlyctis* | 0 | 0 | 0 | 0 | 0.01 | 0 | 0 | 7.40 | 0 |
| *Lichtheimia* | 0.03 | 3.90 | 0.03 | 0 | 0.02 | 0.06 | 0 | 0 | 0 |
| *Solanum* | 0.01 | 0 | 0 | 0 | 0 | 0 | 2.27 | 0.40 | 1.21 |
| *Candida-Lodderomyces_clade* | 0.07 | 0.09 | 0.06 | 0 | 0.03 | 3.55 | 0 | 0 | 0.01 |
| *Cystobasidium* | 0.46 | 0.09 | 0.02 | 0 | 1.36 | 0.03 | 0 | 1.47 | 0.03 |
| *Mariannaea* | 0.01 | 0.22 | 2.63 | 0.01 | 0.03 | 0.02 | 0.05 | 0.27 | 0.08 |
| *Vermiconia* | 0.01 | 0.20 | 0.13 | 0.12 | 0.02 | 0.63 | 0.09 | 1.04 | 0.14 |
| *Peloribates* | 0 | 0 | 0 | 0 | 0 | 0 | 2.10 | 0 | 0 |
| *Coniochaeta* | 0.02 | 0.07 | 0.04 | 0 | 0.29 | 1.08 | 0 | 0.01 | 0.01 |
| *Glycine* | 0 | 0 | 0 | 0 | 0 | 0.01 | 0.21 | 0.14 | 1.10 |
| Other | 3.00 | 3.05 | 2.43 | 0.51 | 1.28 | 3.78 | 1.96 | 4.64 | 2.63 |
